# Super-resolution imaging of RAD51 and DMC1 in DNA repair foci reveals dynamic distribution patterns in meiotic prophase

**DOI:** 10.1101/2020.01.06.895680

**Authors:** Johan A Slotman, Maarten W Paul, Fabrizia Carofiglio, H Martijn de Gruiter, Tessa Vergroesen, Wiggert A van Cappellen, Adriaan B Houtsmuller, Willy M Baarends

## Abstract

The recombinase RAD51, and its meiosis-specific paralog DMC1 localize at DNA double-strand break (DSB) repair sites in meiotic prophase nuclei. While both proteins are required during meiotic homologous recombination, their spatial organization during meiotic DSB repair is not fully understood. Using super-resolution microscopy on mouse spermatocyte nuclei, we aimed to define their relative position at DSB foci, and how these vary in time. We show that a large fraction of meiotic DSB repair foci (38%) contained a single RAD51 cluster and a single DMC1 cluster (D1R1 configuration) that were partially overlapping (average center-center distance around 70 nm). The majority of the rest of the foci had a similar combination of a major RAD51 and DMC1 cluster, but in combination with additional clusters (D2R1, D1R2, D2R2, or DxRy configuration) at an average distance of around 250 nm. As prophase progressed, less D1R1 and more D2R1 foci were observed, where the RAD51 cluster in the D2R1 foci elongated and gradually oriented towards the distant DMC1 cluster. This correlated with more frequently observed RAD51 bridges between the two DMC1 clusters. D1R2 foci frequency was more constant, and the single DMC1 cluster did not elongate, but was observed more frequently in between the two RAD51 clusters in early stages. D2R2 foci were rare (<10%) and nearest neighbour analyses also did not reveal pair formation between D1R1 foci. In the absence of the transverse filament of the synaptonemal complex (connecting the chromosomal axes of homologs), early configurations were more prominent, and RAD51 elongation occurred only transiently. This in-depth analysis of single cell landscapes of RAD51 and DMC1 accumulation patterns at DSB repair sites at super-resolution thus revealed the variability of foci composition, and defined functional consensus configurations that change over time.

**AUTHOR SUMMARY:** Meiosis is a specific type of cell division that is central to sperm and egg formation in sexual reproduction. It forms cells with a single copy of each chromosome, instead of the two copies that are normally present. In meiotic prophase, homologous chromosomes must connect to each other, to be correctly distributed between the daughter cells. This involves the formation and repair of double-strand breaks in the DNA. Here we used super-resolution microscopy to elucidate the localization patterns of two important DNA repair proteins: RAD51 and DMC1. We found that repair sites most often contain a single large cluster of both proteins, with or without one additional smaller cluster of either protein. RAD51 protein clusters displayed lengthening as meiotic prophase progressed. When chromosome pairing was disturbed, we observed changes in the dynamics of protein accumulation patterns, indicating that they actually correspond to certain repair intermediates changing in relative frequency of occurrence. These analyses of single meiotic DNA repair foci reveal the biological variability in protein accumulation patterns, and the localization of RAD51 and DMC1 relative to each other, thereby contributing to our understanding of the molecular basis of meiotic homologous recombination.

## INTRODUCTION

During meiosis, correct homologous chromosome pairing and separation requires the repair of programmed, meiosis-specific, DNA double-strand breaks (DSBs), induced by a meiosis-specific topoisomerase type II-like complex (1–3), in species ranging from yeast to mammals. The machinery that generates and repairs the DSBs is meiosis-specific, but contains many proteins that also function in homologous recombination (HR) repair of DSBs in somatic cells (reviewed in (4)). In somatic HR (mainly active during S or G2 phase), the DNA of DSBs is resected, resulting in the formation of two 3’single-strand (ss) DNA ends, coated by the ssDNA binding protein complex RPA. Subsequently, RPA is replaced by the recombinase RAD51. This enzyme forms a protein filament on the DNA and is capable of mediating strand invasion and strand displacement (D-loop formation) (5). This allows subsequent steps in repair, involving recovery of the missing information from the intact sister chromatid.

Meiotic DSB ends are also resected, but in addition to RPA, meiosis-specific ssDNA binding proteins also associate with the processed ssDNA ends (6, 7). RPA is then displaced by the canonical recombinase RAD51, and its meiosis-specific paralog DMC1 (53.6% amino acid identity to RAD51 in mouse) (8, 9). The two recombinases appear to colocalize in mouse spermatocytes and oocytes when imaged with standard microscopy techniques (10, 11). In *A. thaliana* atRAD51 and atDMC1 have been detected as paired foci, indicating that each of the two DSB ends may be coated by a different recombinase (12). However, recent super-resolution imaging in *S. cerevisiae* has indicated that multiple small DMC1 and RAD51 filaments may accumulate on both ends of a meiotic DSB, and paired co-foci were observed at lower resolution (13). Mouse spermatocytes are very suitable for immunocytology, due to their relatively large size, and well organized patterns of chromosomal axes, that can be used to substage meiotic prophase, using antibodies targeting meiosis-specific chromosomal axis proteins such as SYCP2 and SYCP3, that form the platform on which the programmed DSBs are processed (14). Here we addressed the nanoscopic localisation of RAD51 and DMC1 during mouse meiotic prophase. First, we assessed the overall distribution of RAD51/DMC1 foci in the nucleus using confocal microscopy. Next, we employed a combination of Structured Illumination Microscopy (SIM) and direct Stochastic Optical Reconstruction Microscopy (dSTORM) in two colours to visualize nanoscopic details of RAD51 and DMC1 foci in mouse meiotic prophase nuclei. We compared the localization pattern of the two recombinases in wild type spermatocytes with spermatocytes lacking the transverse filament protein SYCP1 (*Sycp1-/-*). In the absence of this core component of the synaptonemal complex homologous chromosomes align but fail to synapse, resulting in the persistence of meiotic DSB repair foci (15).

Our results show that most repair foci contain single RAD51 and DMC1 clusters that are in close proximity to each other, with or without one much smaller additional RAD51 or DMC1 cluster at larger distance. As prophase progresses, configurations become more complex, and the major domain elongates, but this is dependent on the presence of SYCP1. One of the possible interpretations of these data may be that D1R1 configurations represent filament formation on one end of a meiotic DSB, and that the distance to the other end is highly variable, precluding frequent observation of co-foci. In addition, the relatively frequent occurrence of the D2R1 and D1R2 configurations indicate that there may be stochastic variations in filament formation and/or in chromatin binding patterns of RAD51 and DMC1. This work is a first step towards unravelling the exact molecular composition of the meiotic recombination machinery in time and space in single cells.

## RESULTS

### Non-random distribution of RAD51-DMC1 foci along axial elements

Previous analyses performed on *s. cerevisiae* meiocytes have indicated non-random occurrence of pairs of RAD51-DMC1 co-foci (13). RAD51 and DMC1 also colocalize in easily discernible repair foci in mouse spermatocytes and oocytes (8, 11) but formation of pairs of such foci has not been described, and is also not immediately evident from the microscopic images that can be obtained (Figure 1A). In mouse, these foci are usually analysed in combination with visualization of the axial/lateral elements of the SC, since it is known that the meiotic DSBs localize along these axes. Previously, non-random distribution of markers of repair foci along the axial elements of specific chromosomes has been shown for late zygotene and pachytene spermatocytes, providing evidence for different levels of crossover interference (16–18), but such analyses have not been performed for earlier stages. To ensure nonbiased quantification of immunosignals we selected foci (using FIJI, see Materials and Methods) that were located on the chromosomal axes of leptotene and zygotene nuclei (examples of selected foci and raw images are shown in Figure 1A, C) and determined the nearest distance between RAD51 and DMC1 foci, as well as the RAD51-RAD51 and DMC1-DMC1 distances (Fig. 1B, D). We counted the numbers of foci (Supplemental Figure S1A), and used these numbers to simulate random distributions of the same number of artificially generated foci along the areas covering the SYCP3 signal for each nucleus as described in Materials and Methods (see examples in Fig. 1A, C). This analysis showed that 80% (leptotene) and 67% (zygotene) of the analysed DMC1 foci on the chromosomal axes had a RAD51 neighbour at a distance shorter than 300nm (For p-values and other statistical parameters see Supplementary Figure 1B), reflecting the overall colocalization. Analyses of DMC1-DMC1 and RAD51-RAD51 distances also revealed a non-random distribution (Figure 1B, Supplementary Figure 1B), whereby distances between 500 and 800 nm occurred more frequently than expected. This could be explained by the fact that DSB foci are generally excluded from specific regions, such as constitutive heterochromatin and near centromeric areas, causing foci to be in closer proximity to each other than expected based on random distribution. However, the rather sharp peaks of RAD51-RAD51 and DMC1-DMC1 nearest neighbour distances around 800nm in zygotene, indicate additional non-random distribution within the DSB-foci positive SC regions.

**Fig 1:**
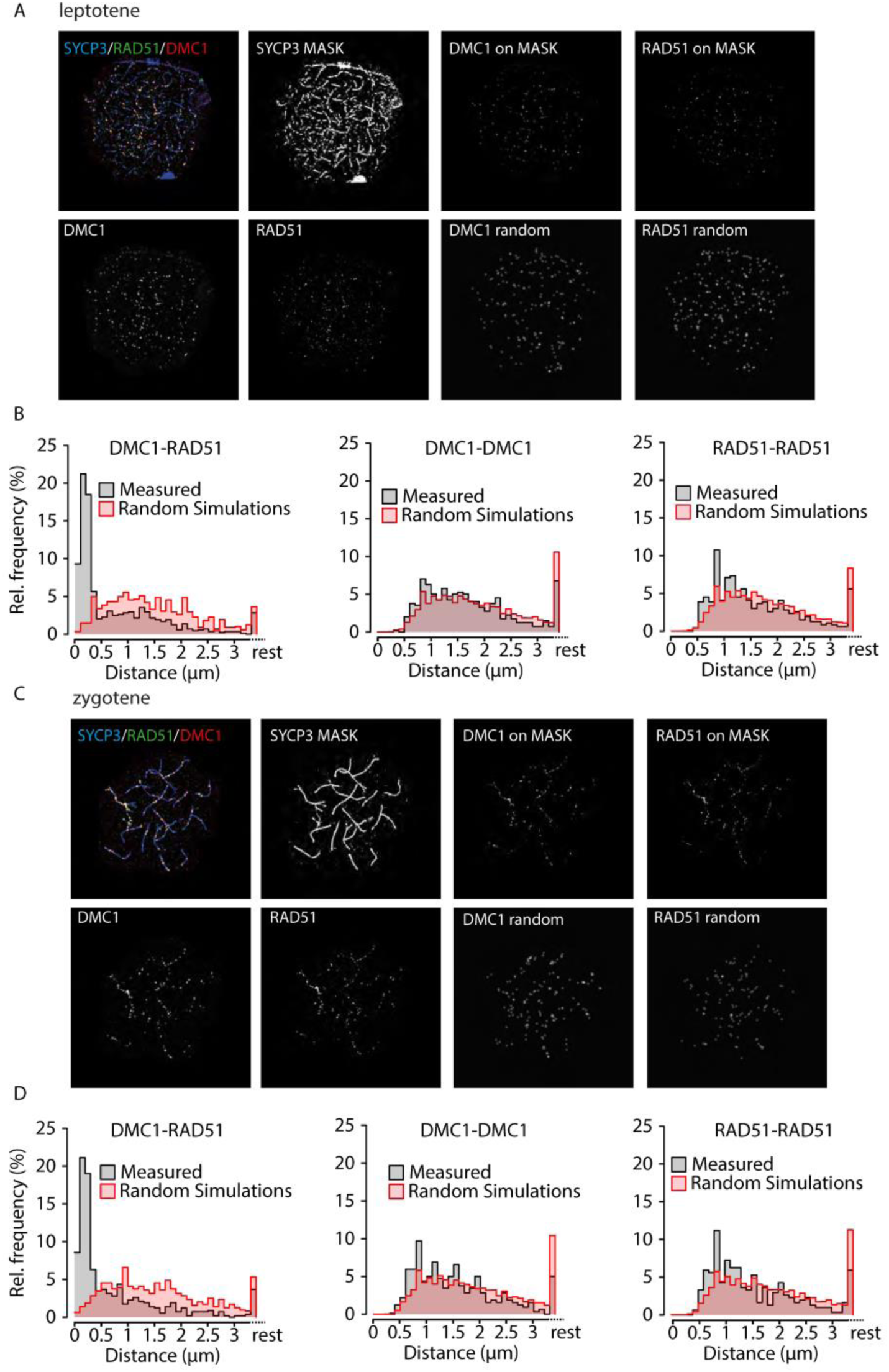
Nearest neighbour analyses of confocal microscopy images of RAD51 and DMC1 foci on the synaptonemal complex axes. A), C) Top left, example confocal image of triple stained leptotene (A) and zygotene (C) nucleus, with primary antibodies for RAD51, DMC1, and SYCP3, and appropriate secondary antibodies conjugated with Alexa 488 (green), Alexa 647 (red), and Alexa 555 (blue), respectively; single DMC1 and RAD51 images are shown in greyscale below; the SYCP3 mask generated as described in Materials and methods is shown to the right of the triple staining; the two top right images show the DMC1 and RAD51 foci that localize on the mask, and below them, the same number of foci randomly distributed on the mask. B), D) Relative frequency distribution of nearest neighbour distances between DMC1 and RAD51 (left) DMC1 and DMC1 (middle) and RAD51 and RAD51 (right) in leptotene (B, n=7 nuclei; 606 DMC1 foci, 712 RAD51 foci) and zygotene (D, n=6 nuclei; 471 DMC1 foci, 462 RAD51 foci) wild type nuclei. Distances were binned in 100nm bins, distances larger than 3.4 µm were labelled as rest. Grey bars, experimental data; red bars, simulated data (see Materials and Methods)

### Composition of meiotic recombination foci revealed by super-resolution imaging

To establish precisely how RAD51 and DMC1 accumulate relative to each other at distances smaller than 300 nm, we visualized RAD51, DMC1, and SYCP3, using SIM and dSTORM, (Figure 2A-E). By utilizing a microscope that combines SIM and dSTORM, we were able to visualise the same field-of-view applying both techniques with the same objective lens (Figure 2A, B). The SIM images were used to visualise synaptonemal complexes (SCs), to be able to identify the substage of meiotic prophase and meiotic DSB foci (also in the SIM image), which were further analysed in images acquired by dSTORM. In DMC1 and RAD51 co-staining experiments, the two proteins displayed distinct localisation patterns, both in SIM and dSTORM images (Figure 2C, D).

**Fig 2:**
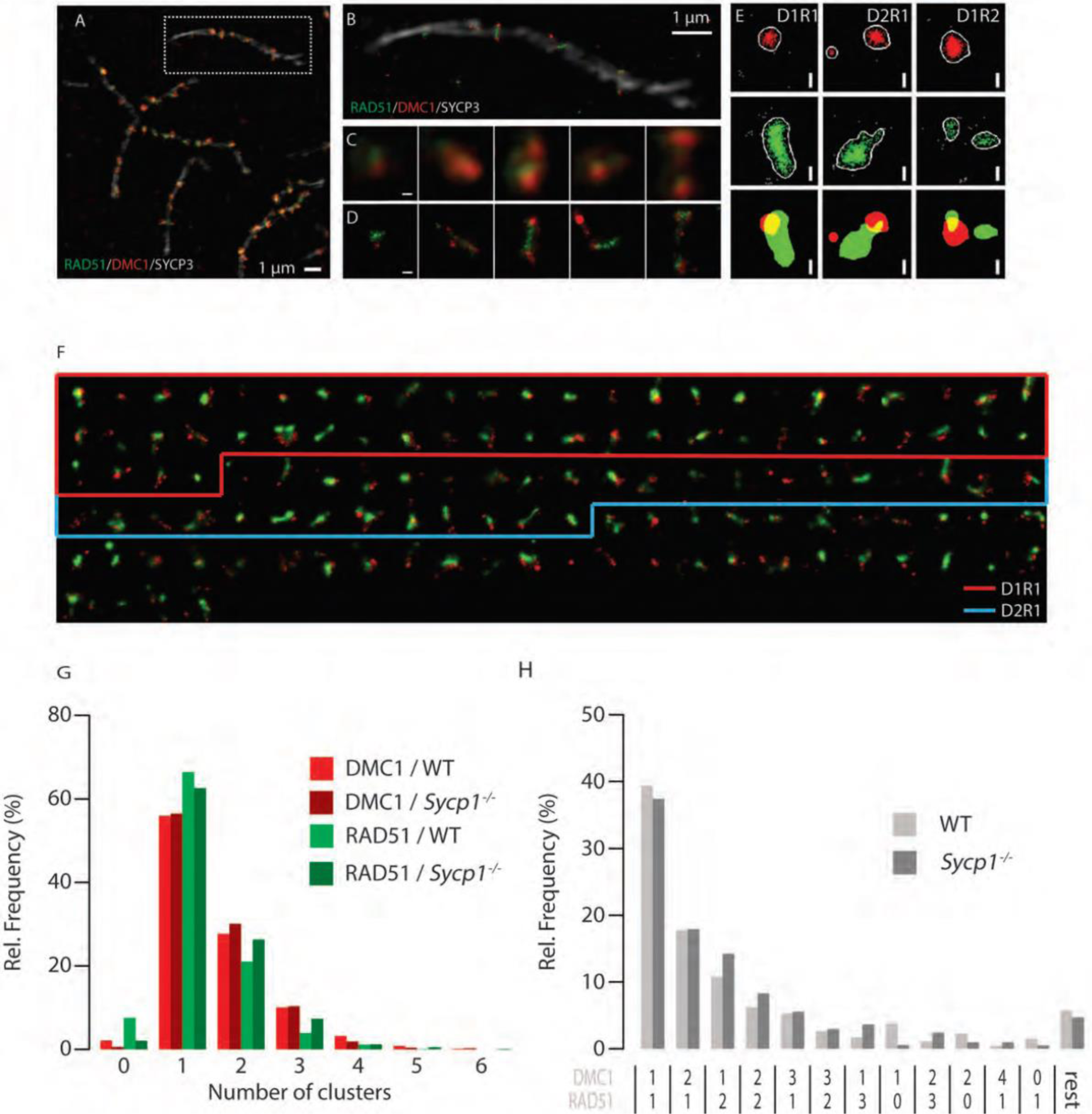
Meiotic DSB foci in super-resolution. A) Cropped region from a SIM image of a spread mouse late zygotene nucleus immunostained with primary antibodies for RAD51, DMC1, and SYCP3, and appropriate secondary antibodies conjugated with Alexa 488 (green), Alexa 647 (red), and Alexa 555 (white), respectively. B) SYCP3 SIM overlayed with RAD51/DMC1 dSTORM images of boxed region in A). C)Close-up of single DSB foci present on the synaptonemal complex shown in A). D)The same foci visualized with dSTORM. E) Single DSB foci of three types (left panels D1R1, middle panels D2R1, right panels D1R2) represented by 2 different visualisation/analysis methods: scatter plot of localisations and merged binary representation of the kernel density estimation. F) Compilation of all ROIs of a single late zygotene nucleus (indicated with an asterisk in Supplemental Figure S2). ROIs are sorted by their DxRy configuration, from most frequent to rare configuration. The boxes indicated the ROIs belonging to the D1R1 (red) and D2R1 (blue) configurations. G) Relative frequency of foci containing indicated number of RAD51 or DMC1 clusters per focus as a percentage of the number of foci per genotype. H) Relative frequency of foci containing the indicated combinations of RAD51 and DMC1 clusters per focus as a percentage of the number of foci per genotype. Combinations that represented less than 1% of the foci in both wild type and Sycp1-/- were grouped in the category referred to as rest. Scale bars 100nm.

A total dataset of 2315 manually selected foci was generated by analysis of 18 nuclei in different meiotic substages, imaged in four independent experiments (Supplemental Figure S2A-C, Supplemental Table S1). The maximum number of foci per nucleus was observed in early zygotene, corresponding well with what we and others have reported previously (11, 19, 20).

### Most foci contain a major domain consisting of one RAD51 and one DMC1 cluster

Many different configurations of RAD51 and DMC1 assemblies can be discerned (Figure 2F). To quantify and categorize the different patterns of RAD51 and DMC1 clusters objectively, we generated binary images and identified specific RAD51 and DMC1 clusters, within the ROIs (600 nm diameter circles) (Figure 2E)(21). We quantified the number of clusters within each ROI and observed that for both RAD51 and DMC1 a single cluster within a ROI was most frequently observed (Figure 2G). Foci with multiple RAD51 or DMC1 clusters were also present, and were somewhat more frequent for DMC1 compared to RAD51 (Figure 2G). Next, we quantified the different RAD51 and DMC1 clustering combinations in our ROIs dataset in order to assess how the two recombinases relate to each other within each ROI. In the distribution of cluster combinations, 68% of the total population of ROIs fell within three specific groups: one DMC1 cluster and one RAD51 cluster (D1R1, 38%), two DMC1 clusters with a single RAD51 cluster (D2R1, 18%), or two RAD51 clusters and one DMC1 cluster (D1R2, 12%) (Figure 2H). Only 6% of the foci contained 2 clusters of each recombinase (D2R2), and all other combinations occurred at lower frequencies.

We also analysed a mouse mutant model in which assembly of the synaptonemal complex (SC) is incomplete due to the absence of the central or transverse filament of the SC (*Sycp1*-/-, 2 animals, two independent experiments, 10 nuclei, 2042 manually selected foci (Supplemental Figure S3, Supplemental Table S1) (15). In spermatocytes from these mice, homologous chromosomes show pairing but no synapsis, and the distances between paired axial elements are larger than between lateral elements in synapsed SCs in the wild type (around 80 nm in wild type and 200 nm in the knockout) (15). In this mutant, leptotene appears normal, and the number of DSB foci observed at this stage is similar to the maximum number observed in wild type spermatocytes, but the failure to synapse disturbs subsequent stages, and prevents completion of meiotic DSB repair ((15); (22–25) and Supplemental Figure S3). Overall, DxRy configurations were present in similar frequencies in wild type and *Sycp1*-/- nuclei, although D1R2 and other configurations with more than one RAD51 cluster were observed somewhat more frequently in the knockout (Figure 2G, H).

Next, we also classified all binary images based on the observed shapes and sizes of clusters. We frequently observed a structure consisting of a relatively large D cluster and a large R cluster with roundish shapes, that partially overlapped. (Figure 3A: “simple”). If the number of D and/or R clusters was large than 1, we also frequently observed this simple structure, and the additional RAD51 and/or DMC1 clusters were then usually smaller than the two main D and R clusters. This simple structure was less frequently observed in zygotene- and pachytene-like *Sycp1-/-* spermatocytes (Figure 3B).

**Figure 3:**
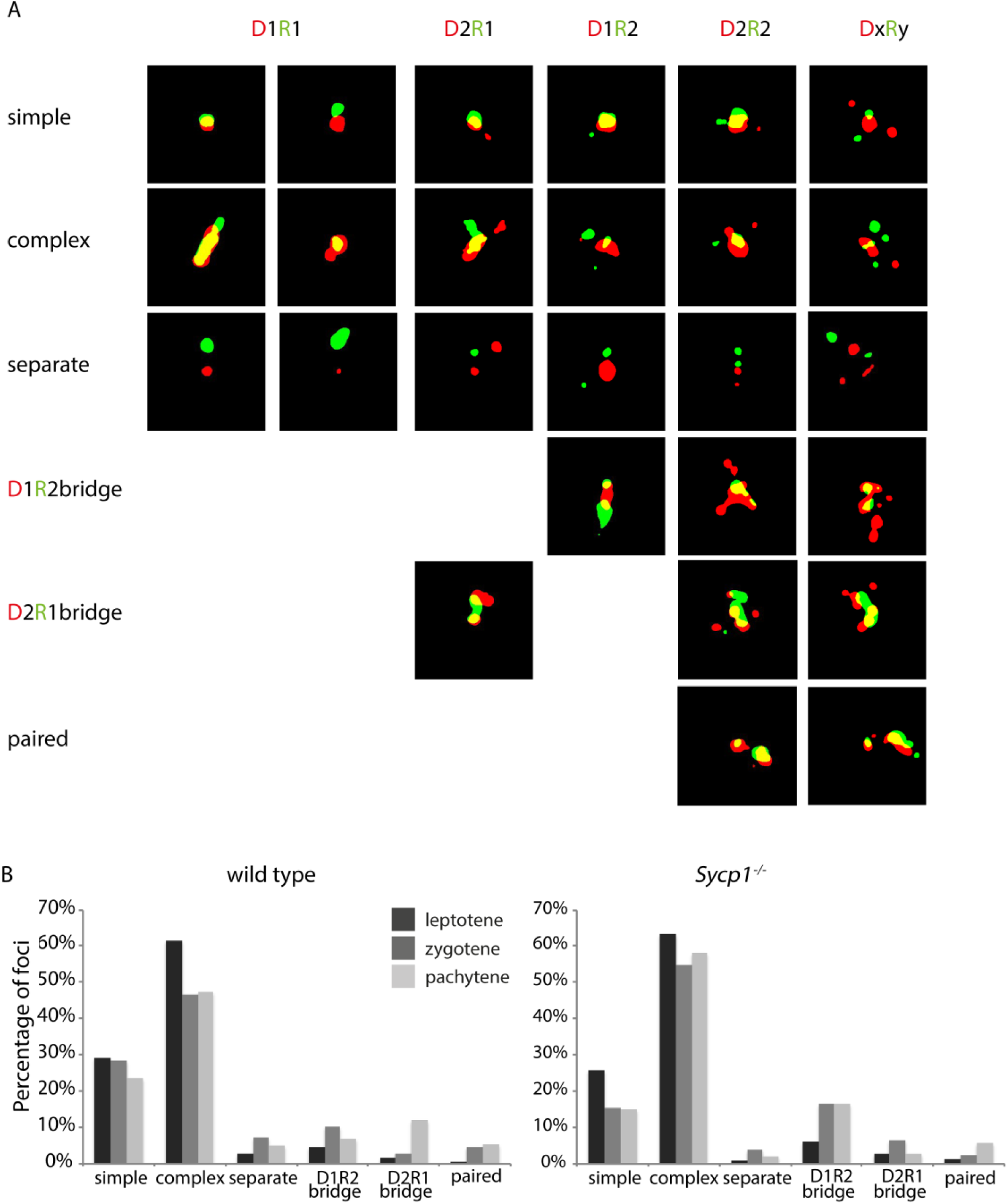
Morphological classification of RAD51-DMC1 configurations. A). All foci were classified as simple, complex, separate, D1R2 bridge, D2R1 bridge, or paired as described in the main text. Examples of each are shown for (from left to right), D1R1 D1R2, D2R1, D2R2, DxRy. B) Relative frequency distributions of the morphological classifications in leptotene (dark grey), zygotene (gray) and pachytene (light gray) of wild type (left) and Sycp1-/- nuclei.

A combination of more complex partially overlapping shapes of a major D and major R domain was also frequently observed in both wild type and *Sycp1-/-* spermatocytes (Figure 3A: “complex”). Again, additional clusters were usually relatively small compared to the two main clusters. Together, these so-called simple and complex foci comprised the majority of all configurations in both wild type and *Sycp1-/-* nuclei. This indicated that the D1R1 foci could actually be representative for a much larger fraction of the DnRn foci if the small additional clusters were considered “satellites”.

A notable structure that was observed for D1R2, D2R1, and ROIs containing more clusters, was termed “bridge” (Figure 3A “bridges”, 13% of all foci in wild type and 19% in *Sycp1-/-*). These contained 2 D clusters that were connected by one R cluster (D2R1 bridge), or the reverse situation (D1R2 bridge), with or without additional clusters. D2R1 bridge frequency increased as prophase progressed in the wild type, but not in the *Sycp1-/-* spermatocytes (Figure 3B). Conversely D1R2 bridges were observed more frequently in zygotene- and pachytene-like *Sycp1-/-* spermatocytes compared to wild type (Figure 3B). Special attention was given to the occurrence of what could be considered as paired configurations; a twin set of overlapping RAD51 and DMC1 clusters (Figure 3A: “paired” and Supplemental Figure S4). These should be mostly represented in the D2R2 subgroup. However, only 34 of the total of 142 D2R2 foci in the wild type have a “paired” appearance (Supplemental Figure S4). The overall frequency of paired configurations increased as prophase progressed in both wild type and *Sypc1-/-* spermatocytes, but never exceeded 6% of the total (Figure 3B).

Finally, a small rather constant fraction of the foci contained only separate RAD51 and DMC1 clusters (Figure 3A: “separate”, and 3B). Given the high relative frequencies of the D1R1, D2R1 and D1R2 configurations in both wild type and *Sypc1-/-* spermatocytes, we investigated these configurations in more detail.

### Temporal analysis of D1R1, D2R1, and D1R2 configurations during meiotic prophase

In wild type nuclei, the D1R1 configuration was the most abundant configuration at leptotene, suggesting that this is an early configuration (Fig. 4A). In the transition to zygotene in the wild type, a reduction of the relative D1R1 configuration frequency was observed, parallel to a 2-fold increase in the relative frequency of D2R1 foci. In contrast, the relative D1R2 frequency remained constant. In *Sycp1-/*- spermatocytes, the absolute and relative D1R1 configuration frequency decreased only transiently in zygotene, and cells that reached a pachytene-like stage displayed D1R1 foci at a frequency that was similar again to what was observed for leptotene nuclei (Fig 4B). The frequency of D2R1 foci remained constant during the different analysed stages of the *Sycp1* knockout (Figure 4B), whereas the D1R2 configuration frequency increased as prophase progressed. Interestingly, this configuration was the only one that appeared to localize preferentially on unsynapsed axes in wild type zygotene nuclei (Figure 4C).

**Fig 4:**
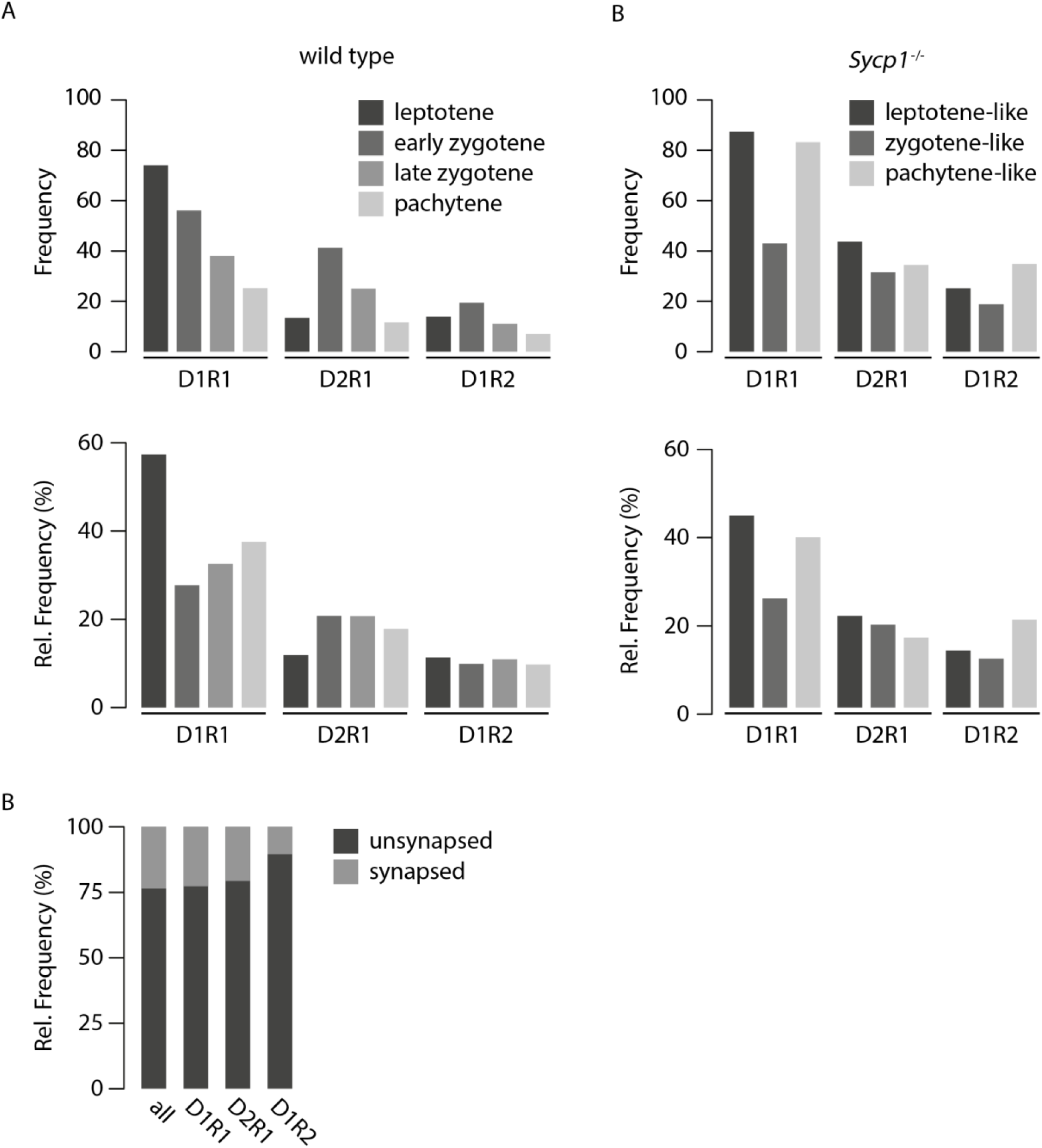
Dynamics of D1R1, D2R1, and D1R2 foci numbers during progression of meiotic prophase in wild type and Sycp1-/- spermatocytes. A) Average frequency (top) and relative frequency (bottom) of D1R1, D2R1, and D1R2 foci per cell per stage for wild type spermatocytes. B) as in A) but for Sycp1-/- spermatocytes. C) Relative frequency (right) of all, D1R1, D2R1 and D1R2 foci on synapsed or unsynapsed synaptonemal complexes at the zygotene stage.

In general, we did not observe any overt specific distribution pattern of the different configurations relative to each other along the SC at the different stages of meiotic prophase (Supplemental Figure S5A).

### Asymmetrical distribution of RAD51 and DMC1 relative to each other in D2R1 and D1R2 configurations

To investigate the spatial organization of protein clusters in the most frequently occurring configurations further (D1R1, D2R1 and D1R2), we determined the center of mass of every cluster in each ROI and measured the distance between the center of RAD51 cluster(s) and DMC1 cluster(s) (Figure 5A, B). Interestingly, minimum distances coherently clustered at approximately 70 nm (wild type/*Sycp1-/-*;68.4±1.2sem/75.8±1.1sem) for all analysed foci configurations in wild type and *Sycp1* knockout nuclei. Thus, almost all foci that contain more than one RAD51 and/or DMC1 cluster, contain at least one RAD51 and one DMC1 cluster in close proximity to each other, with an average distance of approximately 70nm (Figure 5A). Since only a single cluster is present for each of the individual recombinases in the D1R1 group, the distribution of the maximum distance was the same as for the minimum distance. Importantly, it completely overlapped with the first peak of the distribution of maximum distances of all configurations, suggesting that all foci with more than one RAD51 and/or DMC1 cluster, also contain RAD51 and DMC1 clusters that are larger (localisations are more spread) or that are spatially more separated from each other, with an average distance of around 300 nm (wild type/*Sycp1-/-*;287.4±2.7sem/308.6±2.8sem) (Figure 5B).

**Fig 5:**
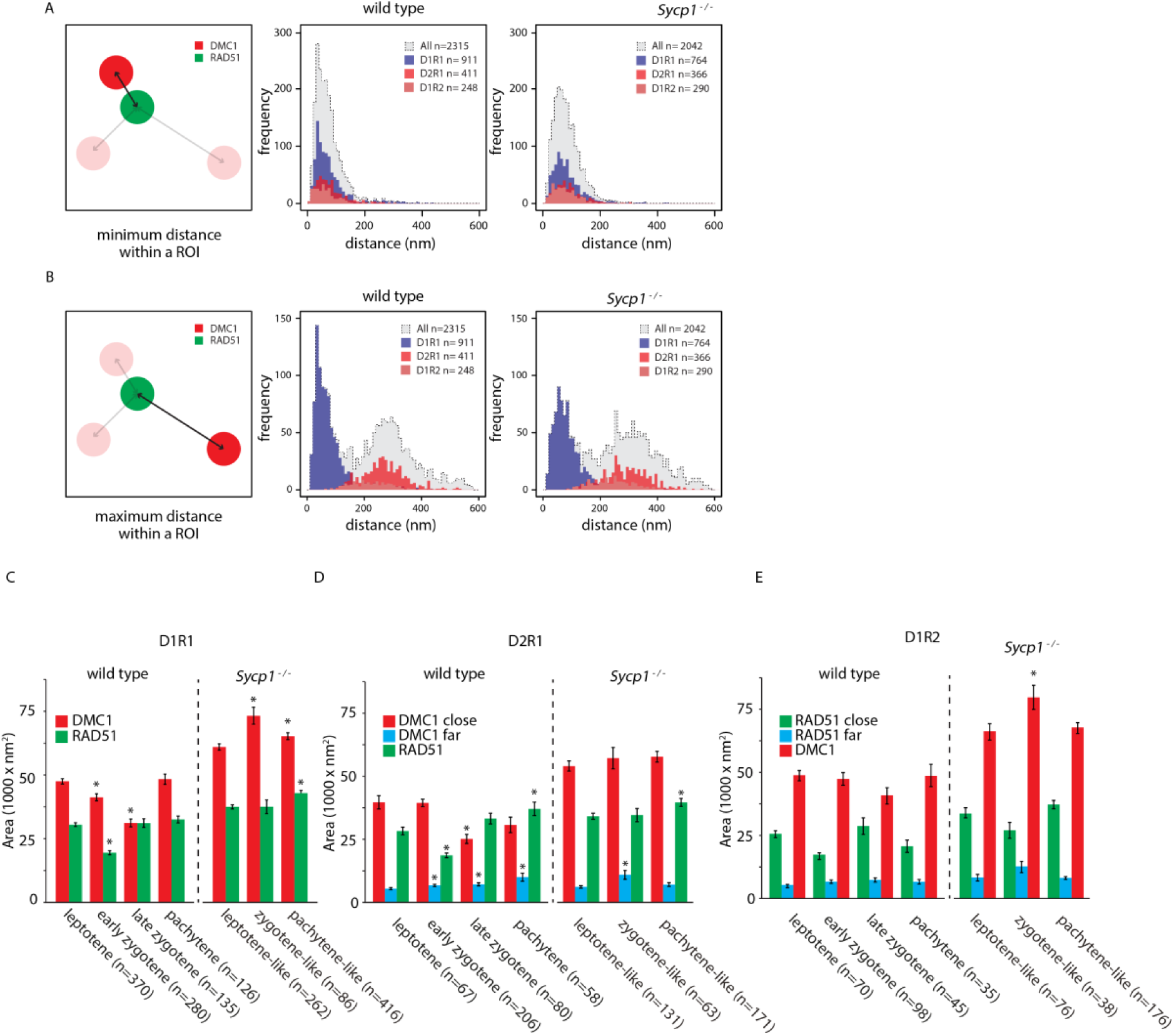
Distances between DMC1 and RAD51 clusters, and area occupancy. A) Distribution of the minimum distances between the center of mass of RAD51 and DMC1 clusters in wild type (middle panel) and Sycp1-/- (right panel) foci. Dashed lines with grey fill represent all foci, the D1R1, D1R2 and D21R1 subgroups are depicted in blue, light red, and red histograms, respectively. B) As in A) but maximum distances are depicted. C) Area of RAD51 and DMC1 clusters in D1R1 subgroup. Error bars indicate SEM, asterisks indicate significant difference compared to leptotene (p<0.05). n indicated number of foci. D) As in C) but area of RAD51 and DMC1 close and far clusters in D2R1 subgroup are shown. E) As in D) but area of RAD51 and DMC1 close and far clusters in D1R2 subgroup are shown. p-values can be found in Supplemental Table S2.

This observation of asymmetry allowed us to define close and far clusters in both the D2R1 and the D1R2 configurations. Interestingly, we observed a large close cluster and a small far cluster irrespective of whether two RAD51 or two DMC1 clusters were present (Fig. 5C-E). Thus, the measured larger distance between the two DMC1 or RAD51 clusters in the D2R1 and D1R2 configurations can be interpreted as more spatial separation. DMC1 area sizes of the close clusters and single clusters are all rather similar in wild type nuclei, and the same holds true for close and single RAD51 clusters. Still, the areas of these large clusters were transiently somewhat decreased in the D1R1 and D2R1 configurations. In addition, the far-DMC1 cluster in the D2R1 displayed a small but gradual increase in size as meiotic prophase progressed. Of note, RAD51 area sizes and DMC1 area sizes did not change during prophase for the D1R2 configuration.

In *Sycp1-/-* spermatocytes, no consistent patterns in area size changes as prophase progressed were apparent (Figure 5C-E).

### Consensus patterns of the spatial organization in D1R1, D2R1 and D1R2 foci

One factor that will contribute to the observed variation in the organization of the individual images is the representation of three-dimensional structures onto a two-dimensional image. To obtain more insight in the actual structure of the three main DxRy configurations, we used alignment by rotation to be able to detect possible consensus patterns in D1R1, D2R1, and D1R2 foci (Figure 6, 7). For the D1R1 group, the DMC1 cluster was used as an anchor point and the RAD51 cluster was used for the rotation. We rotated the structures so that the center of the RAD51 cluster was aligned along the vertical axis above the DMC1 cluster. Then we generated a single fused image of all aligned foci, pooled from the nuclei that were at a specific stage of meiotic prophase. We observed that the RAD51 and DMC1 cluster partially overlap, but that the degree of overlap decreases while meiosis progresses, while the distance between the two clusters increases (Fig. 6A,C). In *Sycp1-/-* D1R1 foci, the degree of overlap was also reduced at the zygotene-like stage, relative to the leptotene-like stage, but increased again at pachytene (Figure 6B). Accordingly, the RAD51-DMC1 distance increases only transiently at the zygotene-like stage (Figure 6C). We observed no differences in distances between clusters within configurations on synapsed versus unsynapsed axes (Supplemental Figure S5B). For D2R1 we used the close DMC1 cluster as anchor, and first rotated the RAD51 cluster along the vertical axis. The resultant locations of the signals of the far-DMC1 cluster were then highly variable at leptotene, but formed a crescent moon-shaped structure around the other two clusters in zygotene and pachytene nuclei (Fig. 6A). As meiotic prophase progresses, the far-DMC1 cluster is more and more localised in a smaller region above the close-DMC1 cluster and the RAD51 cluster, showing that a relatively large fraction of the D2R1 foci has a DMC1-RAD51-DMC1 type of structure. We then aligned the two DMC1 clusters and assessed the RAD51 location relative to the two DMC1 clusters by quantifying the relative number of RAD51 localisations present in four quarters (above, below, left and right) of the image, relative to the close-DMC1 cluster. As expected, based on the results of the rotation with the far-DMC1 cluster, the highest percentage of the RAD51 signal was observed between the two DMC1 clusters, and more signal accumulated in the upper part of that quadrant as prophase progressed (Fig. 6A). In agreement with this observation, the center of mass of the RAD51 cluster seemed to be extending away from the closest DMC1 anchor cluster as cells progressed from zygotene to pachytene (Figure 6D). The mean distance between the two DMC1 clusters, and between the RAD51 and the far-DMC1 cluster in the D2R1 decreased as prophase progressed, but increased again in pachytene (Figure 6E, F). Overall, the consensus patterns in *Sycp1-/-* spermatocytes were similar, but the configurations were more variable (Figure 6B-F). For example, the directionality of RAD51 towards the far DMC1 cluster was clear at the zygotene-like stage, but lost at pachytene-like. Furthermore, in the analyses of the distances between the clusters of the D2R1 configurations, the distance between the close-DMC1 cluster and RAD51 initially appeared larger compared to the wild type, and increased more when cells developed from leptotene to zygotene, but in pachytene-like *Sycp1-/-* spermatocytes, the distance was similar to what was observed at leptotene. The distance of the far-DMC1 cluster to RAD51 or to the close-DMC1 cluster was large at all stages, in contrast to the reduction observed during zygotene in the wild type (Figure 6B, E-F).

**Figure 6:**
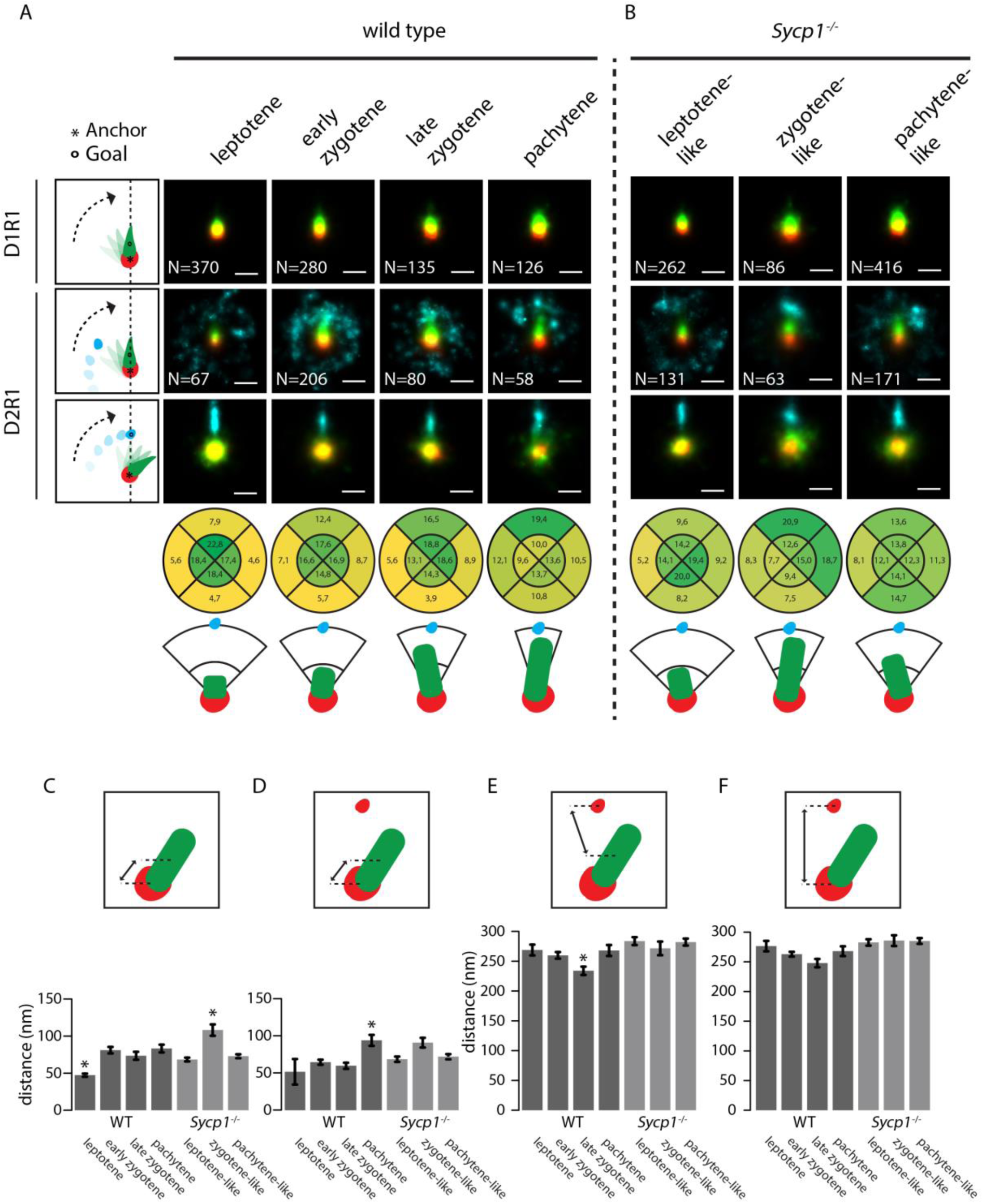
Consensus patterns of D1R1 and D2R1 during meiotic prophase in wild type and Sycp1-/- spermatocytes. Summed images of all rotated and aligned foci within the D1R1 and D2R1 group in wild type (A) and Sycp1-/- (B) per stage. Images were rotated as indicated by schematic drawings to the left of each row, whereby the anchor (*) indicates the cluster that is centred, and the goal (o) the cluster that is rotated to align along the axis. Underneath the lowest D2R1 row, the percentage of localisations for the RAD51 cluster in each indicated quadrant area is shown for each stage for the rotation whereby the close-DMC1 is used as anchor and the far-DMC1 as goal. A schematic interpretation of the results of the rotations is also shown. (C-F) Mean distances between the indicated clusters per stage in wild type and Sycp1-/- spermatocytes. Error bars indicate SEM. Asterisks indicate significant difference compared to all other stages (p<0.05). Scale bars represent 100nm. p-values can be found in Supplemental Table S2.

**Figure 7:**
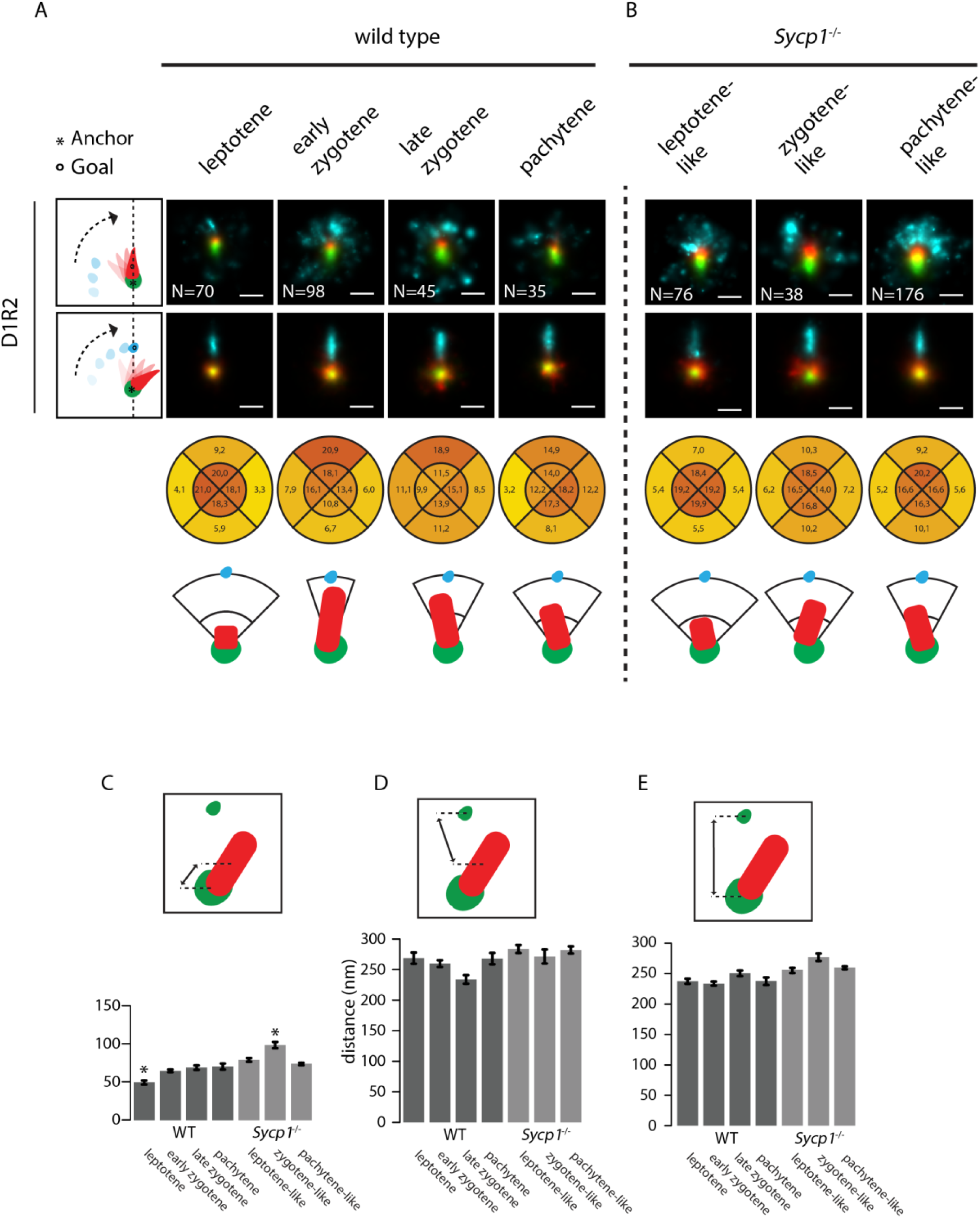
Consensus patterns of D1R2 during meiotic prophase in wild type and Sycp1-/- spermatocytes. Summed images of all rotated and aligned foci within the D1R2 group in wild type (A) and Sycp1-/- (B) per stage. Images were rotated as indicated by schematic drawings to the left of each row, whereby the anchor (*) indicates the cluster that is centred, and the goal (o) the cluster that is rotated to align along the axis. Underneath the lowest D1R2 row, the percentage of localizations for the DMC1 cluster in each indicated quadrant area is shown for each stage for the rotation whereby the close-RAD51 is used as anchor and the far-RAD51 as goal. A schematic interpretation of the results of the rotations is also shown. (C-E) Mean distances between the indicated clusters per stage in wild type and Sycp1-/- spermatocytes. Error bars indicate SEM. Asterisks indicate significant difference compared to all other stages (p<0.05). Scale bars represent 100nm. p-values can be found in Supplemental Table S2.

Finally, we performed the same rotation experiments for the D1R2 configuration. Interestingly, the overall organization of this configuration appeared very similar to the D2R1, including distances between clusters (Compare Fig. 6 to Fig. 7). However, in contrast to the most clear DMC1-RAD51-DMC1 organization of the D2R1 occurring in pachytene, already in early zygotene the single DMC1 cluster of D1R2 was most clearly located between the two RAD51 clusters (Figure 7A), and the DMC1 distance to the close RAD51 was already maximal at early zygotene. No significant change in the distance to the far RAD51 cluster, or between the RAD51 clusters was observed (Figure 7C-E). This corresponds well to the early versus late appearance of the D1R2 and D2R1 bridged structures, respectively (Figure 3B). In *Sycp1-/-* spermatocytes, DMC1 localized more clearly in between the two RAD51 domains, and this was maintained in the pachytene-like nuclei. However, the signal accumulation in the summed rotated images extended less far in the direction of the far RAD51 cluster compared to the wild type (Figure 7A, B). The increase in distance between the DMC1 and close RAD51 cluster was observed only transiently, in zygotene (Figure 7C).

### Three-dimensional simulations of the D2R1 configuration

Next, we simulated a 3D model of D2R1 configurations as described in Materials and Methods (Figure 8A). We analysed the simulated data (discarding the z information) in the same way as the experimental data. Interestingly, around 15% of the simulated D2R1 configurations in a three-dimensional space are represented as D1R1 in the two-dimensional representations, and also a small fraction of D3R1 and D2R2 configurations were observed for simulated D2R1s. This is most likely caused by situations whereby spurious detections rise just above the background, resulting in detection of an additional cluster. We performed rotation and alignment on the simulated D2R1 configurations in the dataset, as described above for the observed real foci. Strikingly, it can be observed that the simulated data fits best to the experimental data set if the degree of freedom for the angle gradually reduces from 132° to 105° and the length of RAD51 gradually increases from 80 to 144 nm going from leptotene to pachytene (Fig. 8C,D). Comparing the simulations to the *Sycp1*-/- D2R1 rotations, it appears that the degree of rotation freedom for the close-DMC1-RAD51 cluster combination relative to the DMC1-DMC1 axis is larger than in wild type at the leptotene-like and pachytene-like stages, but actually more restricted in the zygotene-like nuclei, for which a maximal rotation angle of 105 and a length of 144 nm fitted best.

**Fig 8:**
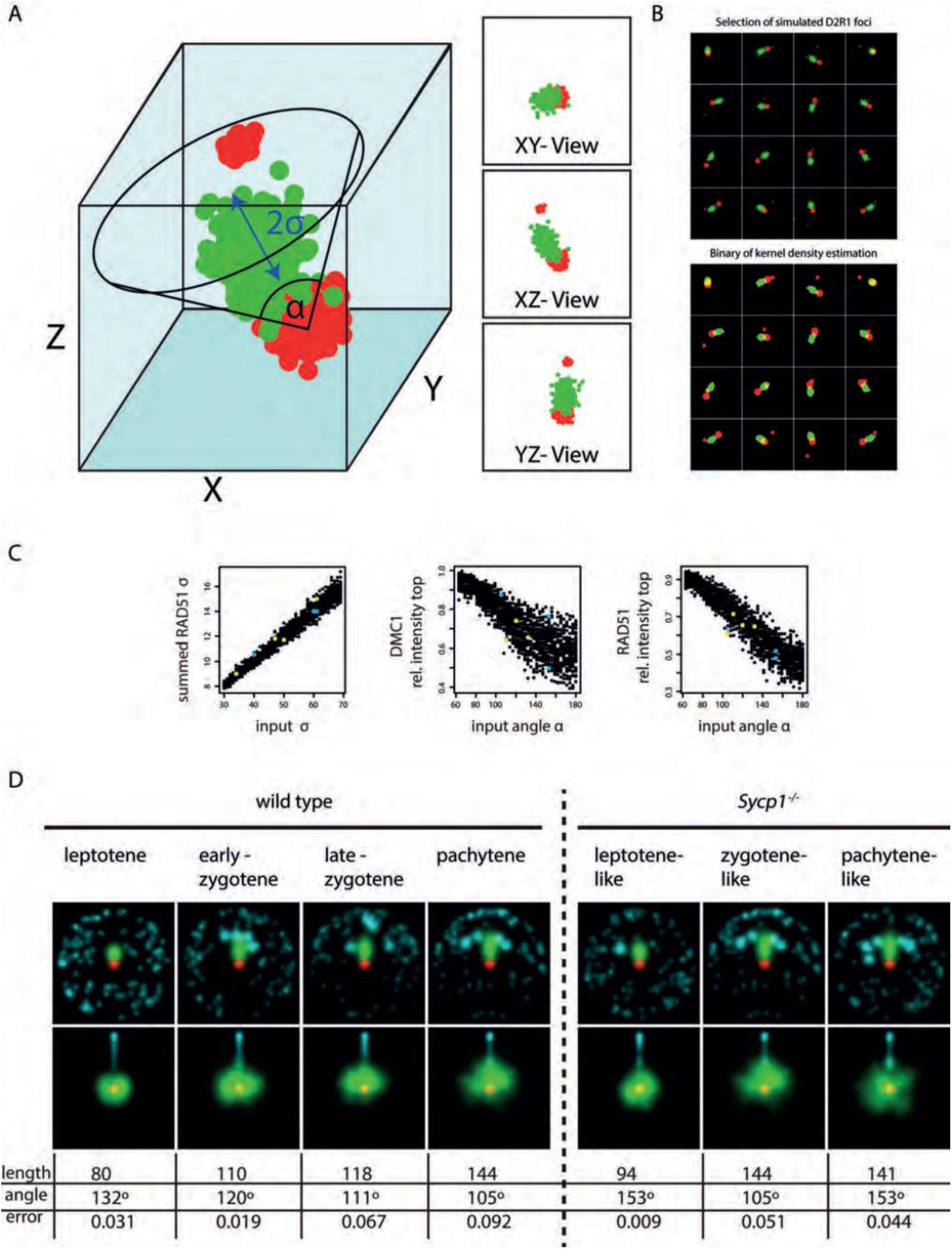
Simulations of D2R1 rotations. A) Model of D2R1 foci in three dimensions, where the alpha indicates the maximum angle relative to the DMC1-DMC1 axis, the sigma the length of the major axis of theRAD51 cluster. B) Selection of simulated foci using one model randomly positioned in space and visualised in two dimensions. C) Measured RAD51 length, RAD51 intensity in the top quadrant and DMC1 intensity in the top half for all simulated foci, whereby each point represents an assembly from 200 aligned foci. Coloured points represent measured values from experimental data from both wild type (yellow) and Sycp1-/- (blue) nuclei at the stages analysed. E) Summed images of simulations that fit best to experimental data, length (full width half maximum: 2.355σ), angle and error are indicated.

## DISCUSSION

We simultaneously determined the localisation of the recombinases RAD51 and DMC1 at nanoscale resolution in more than 4000 DSB foci in 18 wild type and 10 *Sycp1*-/- spermatocytes. We distinguished early, intermediate and late stages of meiotic prophase by co-staining of the synaptonemal complex. Together, this allowed us to reconstruct generalised RAD51 and DMC1 distribution patterns within repair foci as progress through meiotic prophase.

RAD51 and DMC1 filaments are expected to form elongated structures, based on super-resolution images of RAD51 in somatic cells (26, 27). The maximal length of the RAD51 and DMC1 clusters in all observed configurations reached an average of around 140 nm in pachytene, based on our simulations, but the maximal length of the most stretched RAD51 or DMC1 clusters was found to be around 250-300 nm. This is comparable to observed elongated RAD51 structures in fixed somatic cells using dSTORM (26). Haas et al., (27) observed an average maximal length of RAD51 clusters of around 160nm, more similar to the average simulated length observed.

The resolution is limited by the sizes of the first and second antibodies, which is expected to add around 20-40 nm in X and Y direction in our 2D images (28, 29). An *in vitro* filament of RAD51 with a length of 100 nm covers approximately 200 bp of ssDNA (30). Given the current estimates of ssDNA track lengths in meiotic recombination (∼500-1000 bp (31)), it seems reasonable that most of what we observe would represent actual binding of the recombinases to ssDNA, however our data also suggests that neither RAD51 or DMC1 cover the entire resected DNA in an fully extended filament. Furthermore, we certainly cannot exclude that some clusters represent (transient) associations with chromatin, or with dsDNA.

### A close association of a large RAD51 and DMC1 cluster as predominant configuration in DSB repair foci

Since the D1R1 configuration was observed most frequently, and similar structures were also the major component in more complex cluster combinations, the D1R1-configuration represents the main form of RAD51 and DMC1 accumulation at DSB foci. The D1R1 configuration may represent asymmetric loading of each recombinase to one of the two ends of the DSB, or represent loading of both on only one end of a DSB. We hardly observed situations that could be considered paired D1R1 configurations, contrary to what might be expected based on observations in yeast (13), and from the symmetric loading of DMC1 observed in ChIP-seq data of meiotic hotspots (32). Therefore, it appears likely that if D1R1 configurations represent a single end of the DSB, the other end would be mostly occupied by other proteins, or the distance to the other (D1R1) end would be larger than 500nm, and highly variable, precluding visible pairing of D1R1 structures. A combination of these two situations may also occur.

### D2R1 and D1R2 represent DSB intermediates with asymmetric loading of RAD51 and DMC1

The similarity of the DMC1 and RAD51 clusters that are closest to each other in D2R1 and D1R2 to the D1R1 configurations in terms of size and proximity, and the decreasing frequency of the latter, together suggest that the D1R1 may evolve into a D2R1 or D1R2 configuration. The additional cluster at longer distance from the main DMC1-RAD51 entity could then result from new loading of DMC1 or RAD51, or from splitting of the respective cluster into two independent clusters that stabilizes at a distance of 200-250 nm.

The maximum area of the far RAD51/DMC1 cluster is more than 10-fold smaller than the areas occupied by the adjacent close DMC1 and RAD51 clusters. So, either the far clusters may be somehow compacted, or represent binding of recombinase to a shorter stretch of (ss)DNA or chromatin. It is interesting to note that in the protist Tetrahymena, it has been suggested that RAD51 filaments are extremely small, forming no visible foci, whereas DMC1 foci are observed and both proteins are required for functional processing of the meiotic DSBs (33). Although this appears to be an example of extremely asymmetric behaviour of RAD51 and DMC1, our current observations suggest that such small filaments of either RAD51 or DMC1 may also form in other eukaryotes.

Similar to the D1R1, the large close DMC1 and RAD51 clusters in D2R1 and D1R2 may represent binding to the same DNA (single-stranded or double stranded) molecule, or to the different ends of the DSB. The fact that the distances of the two close clusters, to the far cluster in D1R2 and D2R1 foci are very similar in these two configurations supports the idea that there is some form of physical coupling between the D1R1 moiety and the additional RAD51 (D1R2) or DMC1 (D2R1) cluster, and also that the D2R1 and D1R2 configuration represent similar chromatin/DNA conformations/repair intermediates. The “bridged” structures that were observed for both D1R2 and D2R1 also support this notion. The D2R1 bridge was observed mainly in pachytene. This structure, as well as its timing are recapitulated by the lengthening of the RAD51 domain, and increased frequency of DMC1-RAD51-DMC1 alignment as prophase progresses in the rotation analyses. D1R2 bridges were found as more early structures, that preferentially locate on unsynapsed chromatin.

### The number and organization of the RAD51 and DMC1 cluster combinations are affected in *Sycp1-/-* spermatocytes

Our high-resolution analyses revealed an increased frequency of D1R1 configurations in the pachytene-like *Sycp1-/-* nuclei compared to zygotene-like nuclei. Recent data indicate that when synapsis is not achieved, feedback mechanisms may act locally to maintain SPO11 activity in unsynapsed regions (34–36), which is in agreement with the increased frequency of early recombinase configurations in late-stage *Sycp1-/-* spermatocytes. We also observed an increased frequency of D2R1 configurations in leptotene-like nuclei, in comparison to the wild type, which can be attributed to the fact that when a true synapsed structure cannot be formed, initial alignment and pairing will be less stable, and cells that should be in zygotene will still appear as leptotene in the *Sycp1-/-* nuclei. The results of the rotation analyses and distance measurements throughout prophase in the knockout indicate that the D2R1 configuration initially appears to form and proceed as normal, but then a destabilization occurs, leading to frequencies of the D2R1 and D1R2 foci at pachytene-like stage that are more similar to those observed in wild type leptotene cells. This also fits well with a clear increase in D1R2 bridges observed in *Sycp1-/-* nuclei. It is tempting to speculate that in the absence of SYCP1, the lack of SC formation favours D1R2 structures, and that this is somehow coupled to reduced D2R1 formation/stability. In addition, the data support the previously reported longer persistence of DSB induction.

### Concluding remarks

Our super-resolution dual colour dSTORM approach allowed direct comparison of the localization of RAD51 and DMC1 relative to each other. We provide the first evidence for the presence of a major structure consisting of a single relatively large cluster of both RAD51 and DMC1 in close proximity to each other in the majority of mouse meiotic DSB repair foci. Additional, usually smaller clusters of either recombinase are often present, and the fact that the total number of nonoverlapping clusters exceeds two in ∼20% of the foci indicates that some clusters represent binding to dsDNA, or chromatin, or background, since maximally two DSB ends are expected to be available for binding within a single ROI. We favour the hypothesis that the D1R1 configuration mostly represents formation of two adjacent filaments of RAD51 and DMC1 on the same molecule. This then automatically suggests that one DSB end is often not bound by the recombinases, or epitopes are hidden due to differential conformations of the two ends, or the two ends are far apart, with a wide variety in distances, precluding visible formation of paired co-foci.

This single-cell, and single repair focus approach revealed that there is enormous variety in the types of structures formed, in a more or less stochastic manner. We suggest that regulatory mechanisms act to stabilize or destabilize certain structures to eventually allow progression of repair using either the sister chromatid or homologous chromosome at each site, depending on local constraints. Configurations that we observe at low frequencies may still be functionally relevant, and further studies will be required to explain the observed structures in terms of actual repair intermediates. These may involve three-dimensional super-resolution imaging of repair proteins in combination with visualization of DNA. In addition, the experimental combination of meiosis-defective knockout mouse models with super-resolution microscopy provides a promising new approach to study the dynamics of mouse meiotic recombination and meiotic defects at the molecular level.

## MATERIALS AND METHODS

### Animals

Two wild type (5-10 weeks old) and two *Sycp1* knockout (12 weeks old) mice (previously described (15)) were killed using CO2/O2. All animal experiments were approved by the local animal experiments committee DEC Consult and animals were maintained under supervision of the Animal Welfare Officer.

### Meiotic spread preparation and immunofluorescence

Spread nuclei for immunocytochemistry and confocal analyses were prepared as described (37). For dSTORM and 3D-SIM analyses the same method was used, but cells were spread on 1.5 thickness high-precision coverslips (170±5 µm), previously coated with 1% poly L-lysine (Sigma). Slides were immunostained with the antibodies described below in 2 experiments to collect images for the nearest neighbour analyses. Coverslips were stained with antibodies mentioned below in six separate staining experiments for dSTORM and 3D-SIM analyses as follows:

- Four experiments to collect the images of the 18 nuclei presented in Supplemental Figure S2
- Two experiments to collect the images of 10 *Sycp1* knockout nuclei presented in Supplemental Figure S3

Before incubation with antibodies, slides or coverslips were washed in PBS (3×10 min), and non-specific sites were blocked with 0.5% w/v BSA and 0.5% w/v milk powder in PBS. Primary antibodies were diluted in 10% w/v BSA in PBS, and incubations were overnight at room temperature in a humid chamber. Subsequently, slides or coverslips were washed (3×10 min) in PBS, blocked in 10% v/v normal swine serum (Sigma) in blocking buffer (supernatant of 5% w/v milk powder in PBS centrifuged at 14,000 rpm for 10 min), and incubated with secondary antibodies in 10% normal swine serum in blocking buffer overnight at room temperature. Finally, slides or coverslips were washed (3×10 min) in PBS (in the dark) and embedded in Vectashield containing DAPI (slides) or immediately used for imaging 3D-SIM and dSTORM.

### Antibodies

For primary antibodies, we used goat antibody anti-SYCP3 (R&D Systems), mouse monoclonal antibody anti-DMC1 (Abcam ab1837), and a previously generated rabbit polyclonal anti-RAD51 (38). For secondary antibodies, we used a donkey anti-rabbit IgG Alexa 488/647, donkey anti-mouse IgG Alexa 488/647, and donkey anti-goat Alexa 555 (Molecular Probes).

### Confocal imaging

Immunostained spreads were imaged using a Zeiss Confocal Laser Scanning Microscope 700. This microscope is equipped with four lasers with wavelengths of 405 nm, 488 nm, 555 nm and 639 nm. All images were made using a 63x objective immersed in oil with a numerical aperture of 1.40 and a pinhole set at 39 µm. The digital offset was set to -2, and the laser power at 2%. The gain was adjusted for each image and channel. The images are all 1024×1024 in size, averaged 4 times.

### Nearest neighbour analysis

The confocal images were analysed to determine the distribution of RAD51 and DMC1 along the synaptonemal complexes by measuring the nearest neighbour distances. Single nuclei were manually segmented, next DMC1 and RAD51 foci were detected with the ImageJ function “Find Maxima”, and a noise tolerance value of 90 (DMC1) and 100 (RAD51). We then created a mask to outline the SYCP3 signals using manual thresholding, and these masks were then projected onto the image of all the maxima to remove all foci outside the selected area. These masks were also used for the projection of the pixels in the random simulations (see below). The coordinates of the remaining maxima were used to calculate the distances between all the maxima. With these distances the nearest neighbour of each maximum was determined, and the distance values were exported to Excel for further analysis. The nearest neighbour distance distributions of the observed DMC1 and RAD51 foci were compared to random distributions of foci on the SC axes, using the Kolmogorov-Smirnov (KS) test. All KS test values were generated using the R function ks.test.

### Random simulation

Simulated images were created by projecting the number of maxima of a nucleus onto a new image within the boundaries of the SYCP3 signal. This created an image with single pixel foci. To correct for the diffraction limited signal of a confocal microscope, the random image was blurred with a Gaussian filter with a sigma value of 0.11 µm. This sigma value is approximately the standard deviation of the confocal microscope (FWHM = 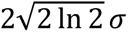 ≈ 2.355 σ (Weisstein, 2002)). Simulated shot noise was added by adding a value of 5 to the entire image, and subsequently adding a random value between +/- the square root of the intensity of each individual pixel. This image was then processed in the same way as the confocal images. 50 random simulations were performed for each nucleus.

### 3D-SIM and dSTORM imaging

Coverslips immunostained as described above were mounted in an Attofluor Cell Chamber (Life Technologies). For drift correction and channel alignment 100nm Gold nanoparticles (Sigma) were added to the sample. To perform dSTORM imaging, an imaging buffer was prepared containing 40mM MEA (Sigma), 0.5mg/ml Glucose Oxidase (Sigma), 40 μg/ml Catalase (Sigma) and 10% w/v Glucose in PBS pH 7.4. Samples were incubated in the imaging buffer during the entire imaging session.

Imaging was performed using a Zeiss Elyra PS1 system. Both 3D-SIM and dSTORM data were acquired using a 100x 1.49NA objective. 488, 561, 642 100mW diode lasers were used to excite the fluorophores together with respectively a BP 495-575 + LP 750, BP 570-650 + LP 750 or LP 655 excitation filter. For 3D-SIM imaging a grating was present in the light path. The grating was modulated in 5 phases and 5 rotations, and multiple z-slices were recorded on an Andor iXon DU 885, 1002×1004 pixel EMCCD camera. dSTORM imaging was done using near-TIRF settings while the images were recorded on Andor iXon DU 897, 512×512 pixel EMCCD camera. At least 10 000 images were acquired at an interval of 33ms for Alexa 647. For Alexa 488 an interval of 50ms was used to compensate for the lower photon yield of the Alexa 488 dye. We used Alexa 488 and Alexa 647 dyes coupled to secondary antibodies to detect respectively RAD51 and DMC1 or vice versa. Using either fluorophore combination, we consistently detected ∼1.5 times more localisation events for RAD51 than DMC1. As expected, we observed more localisations for Alexa 647 compared to Alexa 488, due to the more suitable photochemical properties for dSTORM of the former (39). We chose the more efficient Alexa 647 dye to detect DMC1, that is either less abundant or less well recognized by the primary antibody compared to RAD51, and the Alexa 488 dye to detect RAD51.

### 3D-SIM and dSTORM image analysis

3D-SIM images were analysed using the algorithm in the ZEN2011 (Carl Zeiss, Jena) software. For dSTORM, individual fluorescent events were localised in the subsequent frames using a 2D Gauss fitting algorithm in the ZEN2011 (Carl Zeiss, Jena) software. Detections in subsequent frames originating from the same fluorophore were grouped. Drift was corrected using 100nm gold nanoparticles (Sigma). The same fiducials were used to align the two colour dSTORM images using an affine alignment. Dual colour dSTORM and triple colour SIM images were aligned, based on the dSTORM and 3D-SIM Alexa 647 images, using a channel alignment algorithm in the ZEN2011 software. All observed foci were manually selected based on the SIM images, and circular regions (radius of 300 nm) around the foci were selected using ImageJ within the Fiji platform (40). For each stage and each genotype, 2-5 nuclei were analysed. Each nucleus can be viewed as a biological replicate when differences between stages are considered, whereas each focus can be considered a biological replicate when the overall properties of the foci are analysed. The single molecule localisations of the individual foci were subsequently imported into R using the RStudio GUI for further analysis (Pau, Oles, Smith, Sklyar and Huber, EBImage: Image processing toolbox for R. v. 2.13 (2013) http://watson.nci.nih.gov/bioc_mirror/packages/2.13/bioc/html/EBImage.html; R Development Core Team, R: A language and environment for statistical computing. R Foundation for Statistical Computing, R Foundation for Statistical Computing, Vienna, Austria, ISBN 3-900051-07-0, http://www.R-project.org.)

Selected foci that were spatially overlapping were excluded if the percentage of overlapping localisations was larger than 25% (21). Also foci containing less than 50 localisations were excluded from further analysis.

### Foci analysis

Single molecule localisation data was used to fit a 2D Kernel Density Estimation (KDE) function (Wand, 2013, KernSmooth: Functions for kernel smoothing for Wand & Jones 2.23-10, http://CRAN.R-project.org/package=KernSmooth). The KDE function estimates the density of localisations at a certain position in the image. The bandwidth of the density estimation was set to the approximate average localisation precision of our data: 20 nm. The 2D KDE gives a normalized density over the image. Because we are interested to determine the absolute density of localisations, the normalized density is multiplied by the number of localisations in the ROI. After fitting a 2D KDE to the data we are able to define objects by applying a threshold. The threshold was set at 5 localisation/pixel, equal to 0.2 localisations/nm2. Very small clusters with an area covering less than 50 pixels were considered background. The resulting binary images were used to determine shape features (center of mass i.e.) (Pau, Oles, Smith, Sklyar and Huber, EBImage: Image processing toolbox for R. v. 2.13 (2013) http://watson.nci.nih.gov/bioc_mirror/packages/2.13/bioc/html/EBImage.html).

Pairwise comparison between the mean values of image features from individual meiotic stages was performed using an independent two sample Student t-test. A p-value below 0.05 was considered a significant difference between the two samples. For alignment by rotation the center of mass was used to center images on the close DMC1 cluster for alignment by rotation. The subsequent localisations were all rotated so that either the far DMC1 or RAD51 center aligned above the (close DMC1) center. All localisations from indicated stages were pooled and rendered as an image using SMoLR (21).

### Simulation

We generated a 3D model of a D2R1 focus consisting of three distinct Gaussian distributions of 3D coordinates. The two DMC1 clusters are represented as globular distributions where the standard deviation (σ) of the Gaussian distribution is equal in x,y and z. RAD51 is represented as an ellipsoid distribution in which the σ of the Gaussian distribution is larger in one dimension. We used the mean number of localisations measured per cluster: 267, 564 and 51 coordinates for RAD51, close DMC1 and far DMC1 respectively. We included 50 randomly distributed background coordinates in the model. The model was organized in such a way that the ‘close’ DMC1 cluster and the RAD51 cluster are physically connected. The far DMC1 cluster was placed randomly at distance of 400 nm from the close DMC1 and the RAD51 cluster localises at a random angle relative to the DMC1-DMC1 axis in a three-dimensional space. We then varied the length of the main axis of the RAD51 cluster (σ) and the maximal angle (α) at which the ‘close’ DMC1-RAD51 cluster combination could be positioned relative to the DMC1-DMC1 axis, and generated datasets of 200 configurations for every combination of σ and α. We fitted the experimental data to the simulations using 3 parameters: the σ of a Gaussian fitted over the RAD51 signal (σ-RAD51), the percentage of DMC1 signal in the top half of the center (α-DMC1) in the rotation where RAD51 is aligned to the top, and the percentage of RAD51 in the top quadrant (α-RAD51) in the rotations where the far DMC1 is aligned to the top. These 3 parameters where measured in both the simulated data and the experimental data (Fig 7B). Using a least mean squares method the simulation which fits the experimental data best was determined.

## ACKNOWLEDGEMENTS

We would like to thank Prof. Dr. J. Anton Grootegoed (Erasmus MC, Rotterdam) for his valuable comments and suggestions to the initial manuscript draft and Sabrah Niesten (Erasmus MC, Rotterdam) for her contribution to the code components of the nearest neighbour analyses.

## AUTHOR CONTRIBUTIONS

Conceptualization, FC, MWP, JAS, ABH, and WMB; Methodology, FC, MWP, JAS, WavC, MdG, TV, and ABH; Software, MWP, JAS, and MdG; Formal Analyses, MWP, TV, and JAS; Investigation, FC, MWP, TV, and JAS; Writing-Original Draft, FC, MWP, and WMB; Writing-Review & Editing, all authors; Visualisation, FC, MWP, JAS, and WMB; Supervision, ABH, and WMB; Funding Acquisition, ABH and WMB

## SUPPORTING INFORMATION

**Supplemental Figure S1:**
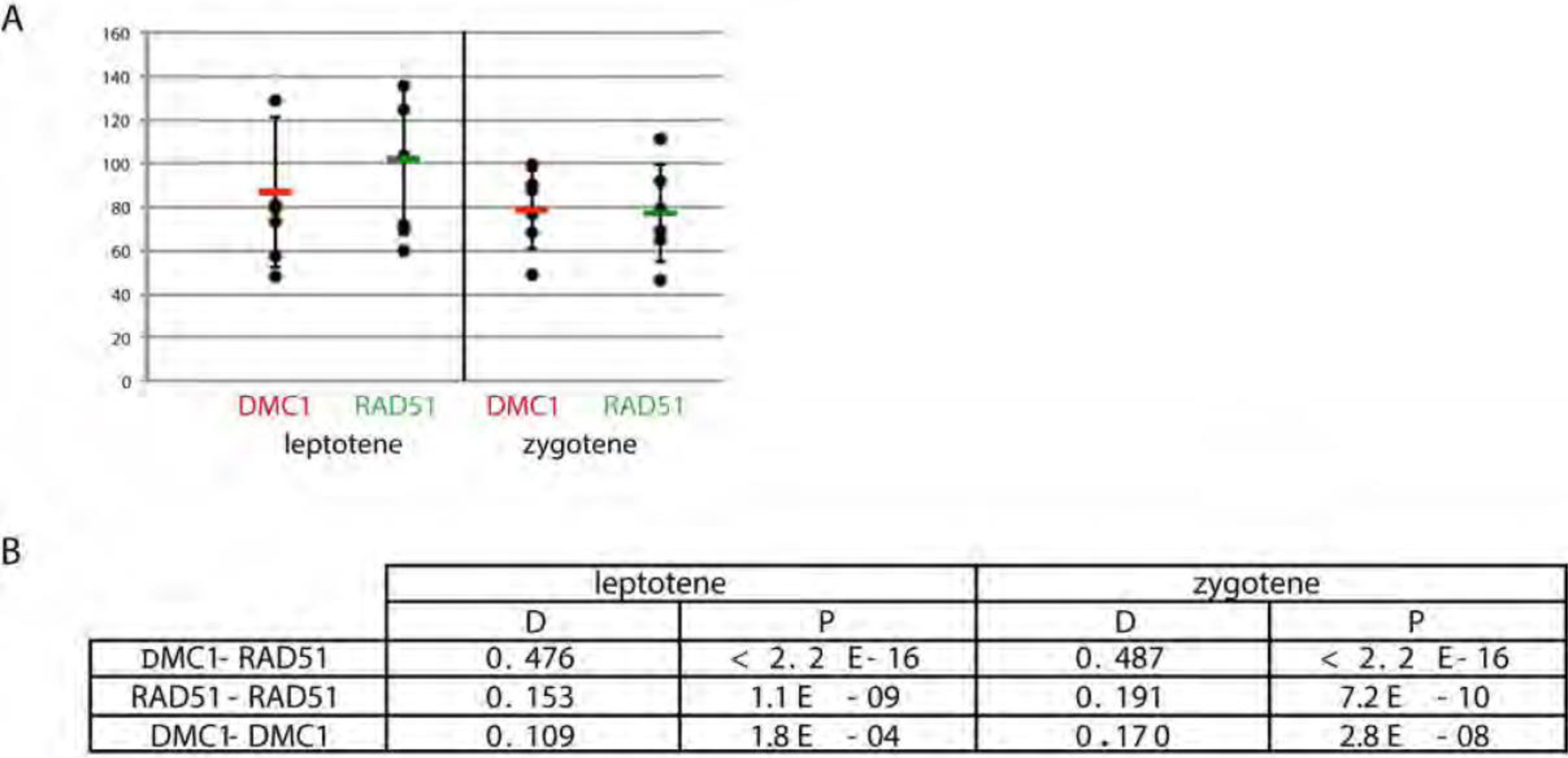
Foci numbers in confocal images used for nearest neighbour distance measurement and statistical analyses A) Foci numbers were determined automatically using FIJI as described in materials and methods. Numbers counted in each individual nucleus are shown. Horizontal bar depicts the average and error bars indicate standard deviation. B) Results from Kolmogorov-Smirnov test comparing nearest neighbour distance distributions between experimental data and simulations. Distance (D) values (largest vertical distance in cumulative frequency histogram of distances) and probability (p) values are shown for the indicated analyses.

**Supplemental Figure S2:**
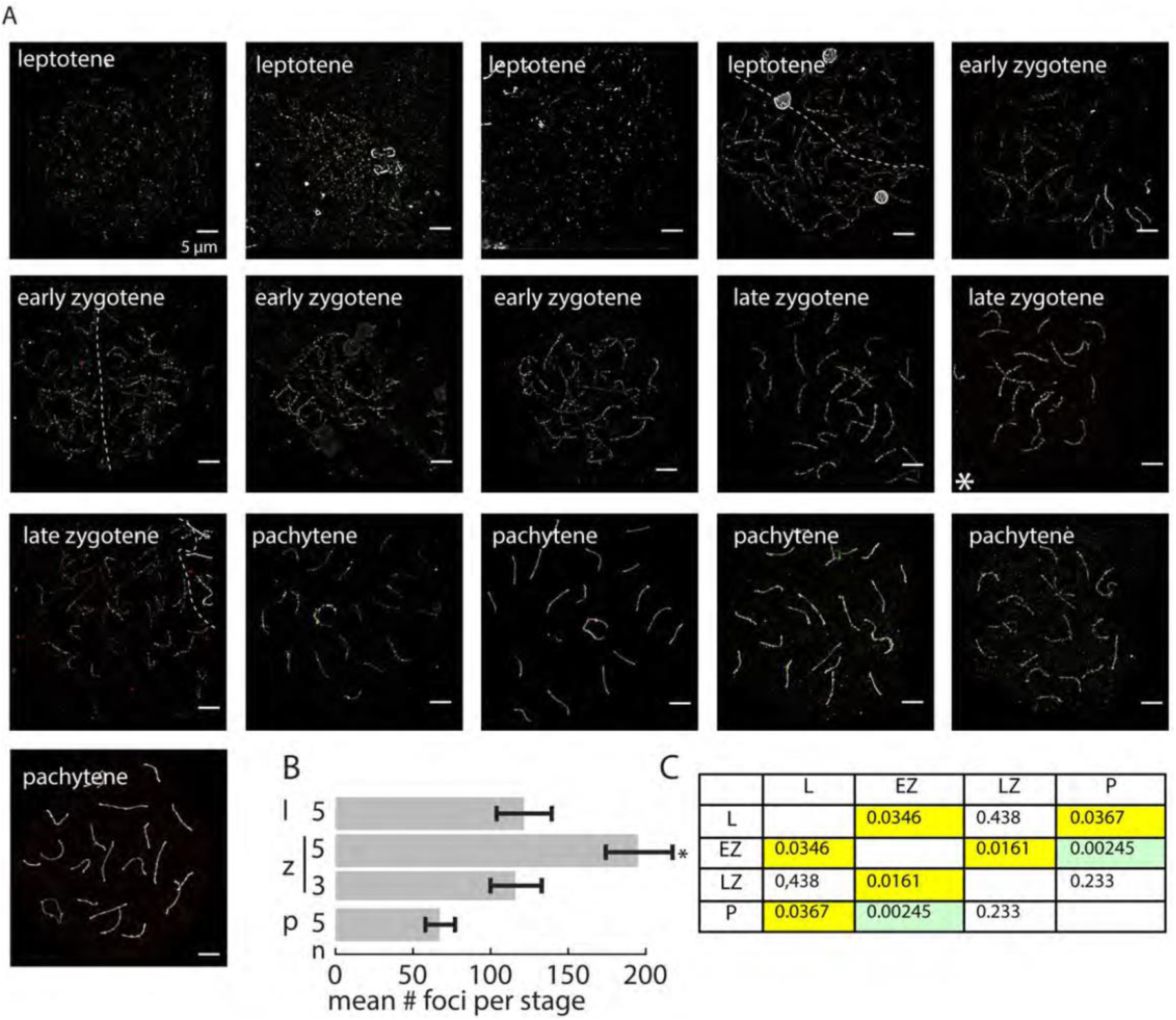
Analysed wild type nuclei (A) 3D-SIM images of the wild type nuclei analysed per stage. Nuclei were immunostained for RAD51 (green), DMC1 (red), and SYCP3 (white). In cases where two nuclei were imaged in the same field of view they are separated by a dashed line. Scale bars represent 5 μm. Asterisk indicates late zygotene nucleus of which foci are shown in Figure 2F (B) Bar graph showing the average number of foci from wild type spermatocyte nuclei that were analysed in dSTORM per stage (leptotene, early/late zygotene, pachytene). The number of analysed nuclei per stage is indicated to the left of each bar. Error bars indicate SEM, asterisk indicate significant difference to all other stages (p<0.05). (C) p-values for foci number comparisons between stages (yellow background; p<0.05, green background p<0.005)

**Supplemental Figure S3:**
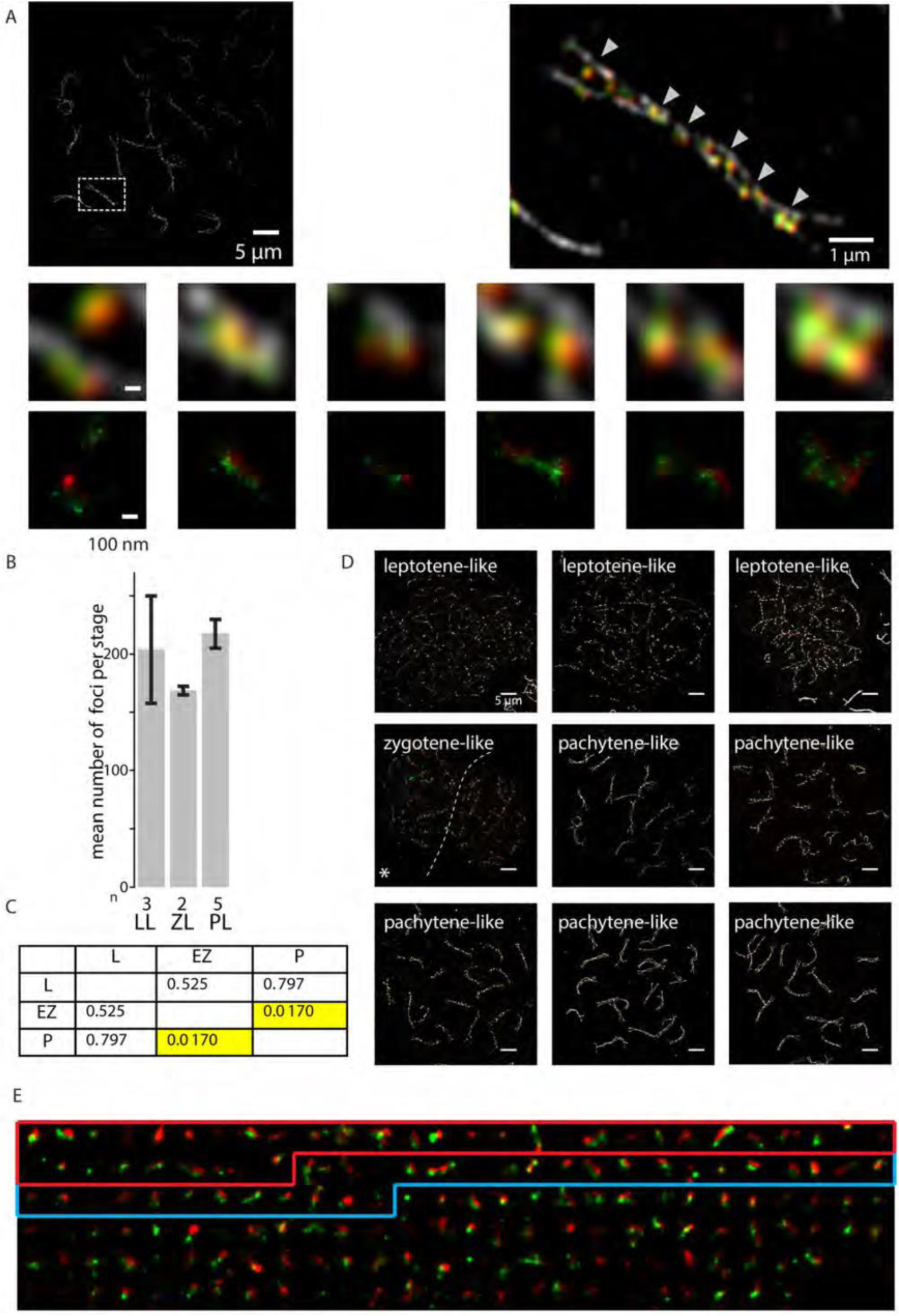
Analysed *Sycp1-/-* nuclei (A) 3D-SIM image of microspread pachytene-like meiotic nucleus from *Sycp1*-/- mouse immunostained with primary antibodies for RAD51, DMC1, and SYCP3, and appropriate secondary antibodies labelled with Alexa 488 (green), Alexa 647 (red), and Alexa 555 (white), respectively. The boxed region is shown to the right and the arrowheads mark regions shown in below. (B) Bar graph showing the average number of foci from wild type spermatocyte nuclei that were analysed in dSTORM per stage (leptotene, early/late zygotene, pachytene). The number of analysed nuclei per stage is indicated underneath each bar. Error bars indicate SEM values. (C) p-values for foci number comparisons between stages (yellow background; p<0.05, green background p<0.005). (D) 3D-SIM images of the *Sycp1-/-* nuclei analysed per stage. Nuclei were immunostained for RAD51 (green), DMC1 (red), and SYCP3 (white). (E) A compilation of all ROIs of the left zygotene-like nucleus, ROIs are sorted by their DxRy configuration, from most frequent to rare configuration. The boxes indicated the ROIs belonging to the D1R1 (red) and D2R1 (blue) configurations. The images are reconstructed with plotted Gaussian distributions proportional to the precision of the individual localisations.

**Supplemental Figure S4:**
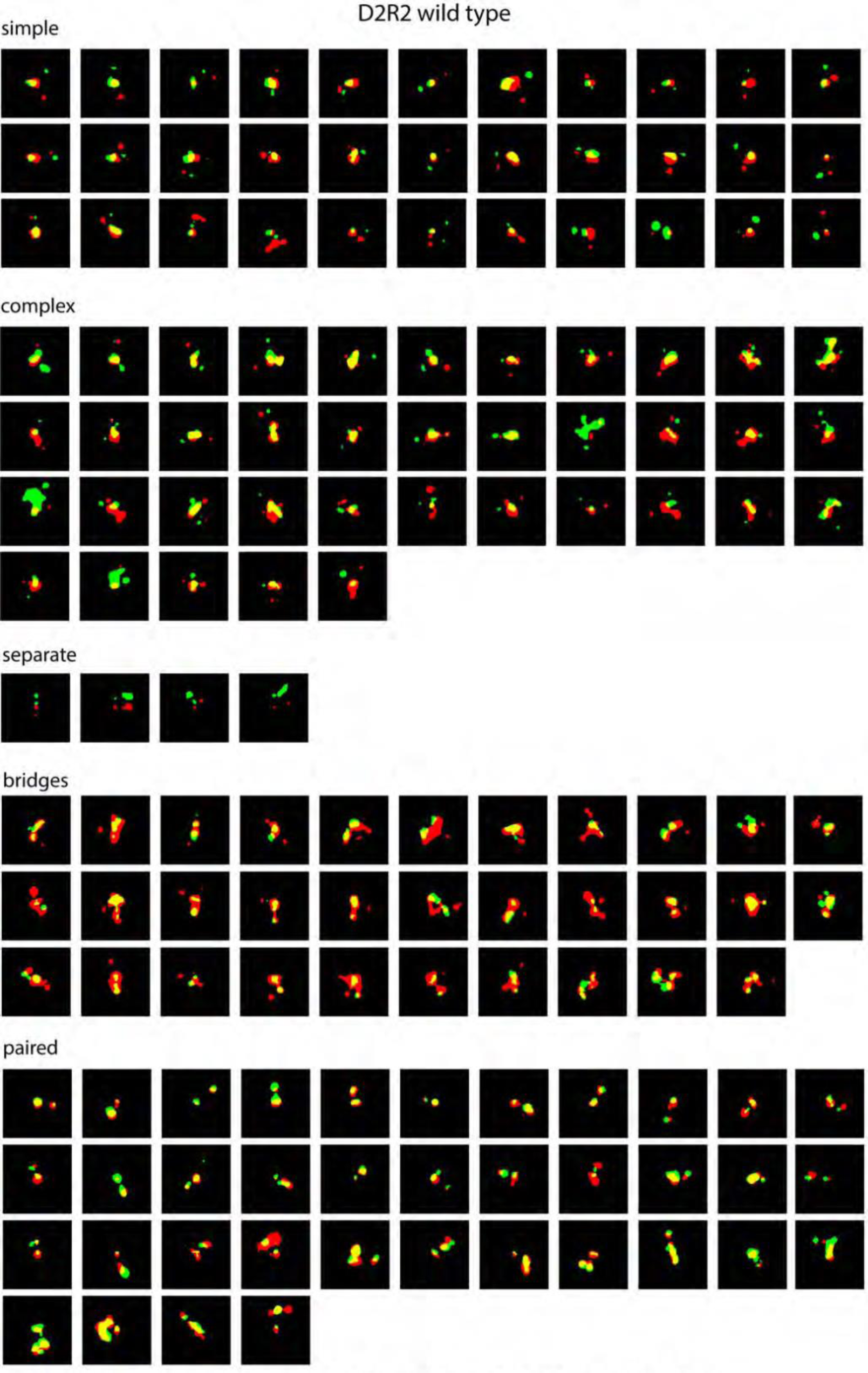
Morphological classification of all wild type D2R2 foci All D2R2 foci are shown, classified as described in the main text

**Supplemental Figure S5:**
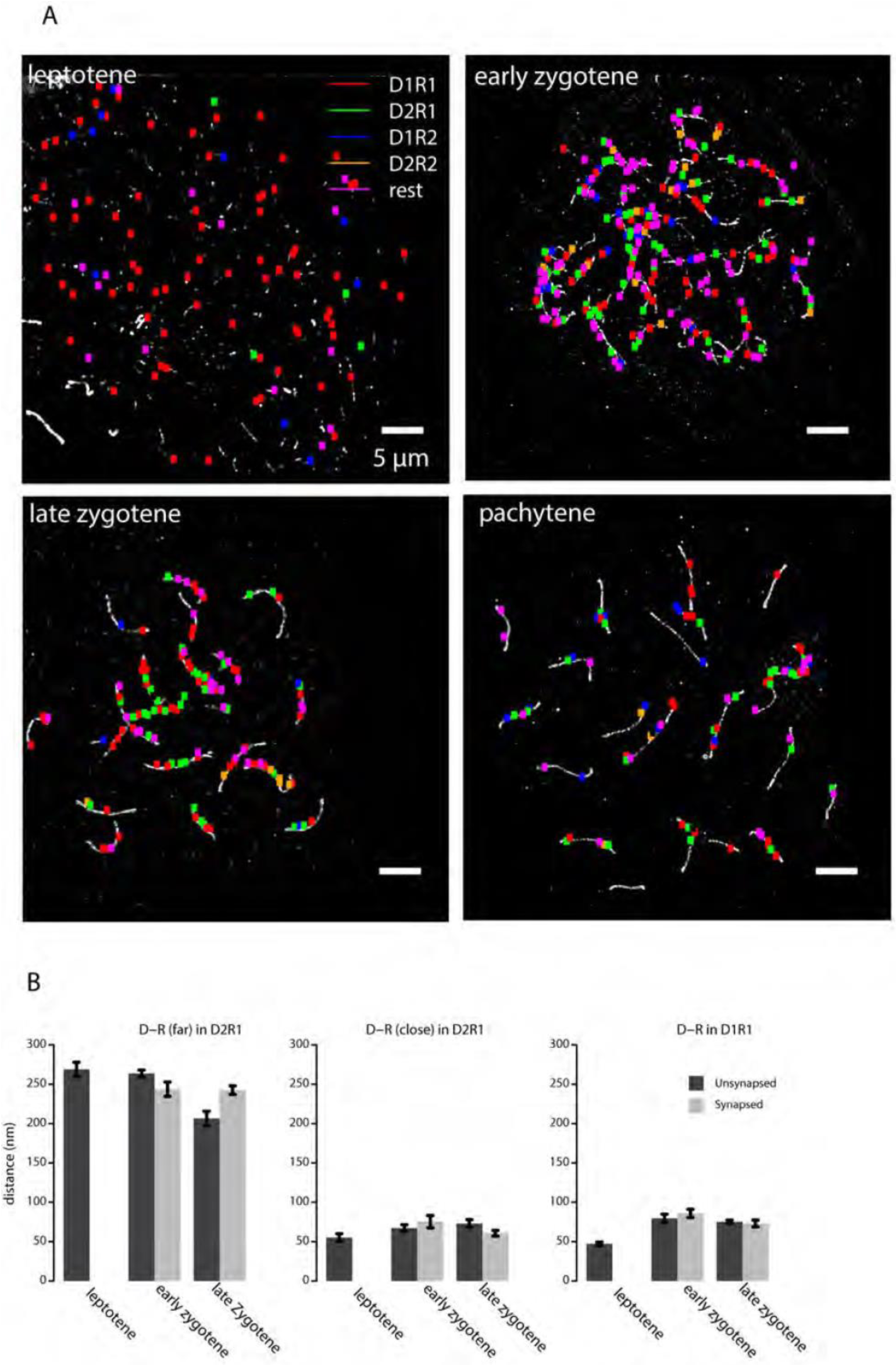
Distribution of different DxRy configurations along the chromosomes of wild type spermatocytes, and analyses of distances between DMC1 and RAD51 clusters on synapsed and unsynapsed axes. (A) The ROIs defined for a wild type leptotene, early zygotene, late zygotene and pachytene nucleus immunostained for RAD51, DMC1 and SYCP3 are superimposed on the SYCP3 SIM image (white). Red ROIs correspond to D1R1, green ROIs correspond to D2R1, blue ROIs to D1R2, yellow ROIs to D2R2 and magenta ROIs to the rest group of configurations. Scale bars indicate 5 µm. (B) Mean distances between the DMC1 and RAD51 clusters in D1R1 and D2R1 configurations per stage in wild type spermatocytes, distributed over synapsed or unsynapsed axes. Error bars indicate SEM.

Supplemental Table S1:

This Excel file contains the data for each focus that was analysed in wild type and *Sycp1-/-* nuclei, as explained in Materials and Methods.

Supplemental Table S2:

This Excel file contains the areas, SD and SEM values used to generate panels C and D of Figure 4. In addition, p values are shown for the different comparisons in Figure 4,6, and 7.

## REFERENCES

1. Baudat F, Manova K, Yuen JP, Jasin M, Keeney S. Chromosome synapsis defects and sexually dimorphic meiotic progression in mice lacking spo11. Molecular cell. 2000;6(5):989–98.

2. Romanienko PJ, Camerini-Otero RD. The mouse spo11 gene is required for meiotic chromosome synapsis. Molecular cell. 2000;6(5):975–87.

3. Robert T, Nore A, Brun C, Maffre C, Crimi B, Bourbon HM, et al. The TopoVIB-Like protein family is required for meiotic DNA double-strand break formation. Science. 2016;351(6276):943–9.

4. Hunter N. Meiotic Recombination: The Essence of Heredity. Cold Spring Harbor perspectives in biology. 2015;7(12).

5. Wright WD, Shah SS, Heyer WD. Homologous recombination and the repair of DNA double-strand breaks. The Journal of biological chemistry. 2018;293(27):10524–35.

6. Inagaki A, Schoenmakers S, Baarends WM. DNA double strand break repair, chromosome synapsis and transcriptional silencing in meiosis. Epigenetics. 2010;5(4):255–66.

7. Ribeiro J, Abby E, Livera G, Martini E. RPA homologs and ssDNA processing during meiotic recombination. Chromosoma. 2016;125(2):265–76.

8. Moens PB, Kolas NK, Tarsounas M, Marcon E, Cohen PE, Spyropoulos B. The time course and chromosomal localization of recombination-related proteins at meiosis in the mouse are compatible with models that can resolve the early DNA-DNA interactions without reciprocal recombination. Journal of cell science. 2002;115(Pt 8):1611–22.

9. Habu T, Taki T, West A, Nishimune Y, Morita T. The mouse and human homologs of DMC1, the yeast meiosis-specific homologous recombination gene, have a common unique form of exon-skipped transcript in meiosis. Nucleic acids research. 1996;24(3):470–7.

10. Moens PB, Chen DJ, Shen Z, Kolas N, Tarsounas M, Heng HHQ. Rad51 immunocytology in rat and mouse spermatocytes and oocytes. Chromosoma. 1997;106:207–15.

11. Tarsounas M, Morita T, Pearlman RE, Moens PB. RAD51 and DMC1 form mixed complexes associated with mouse meiotic chromosome cores and synaptonemal complexes. The Journal of cell biology. 1999;147(2):207–20.

12. Kurzbauer MT, Uanschou C, Chen D, Schlogelhofer P. The recombinases DMC1 and RAD51 are functionally and spatially separated during meiosis in Arabidopsis. Plant Cell. 2012;24(5):2058–70.

13. Brown MS, Grubb J, Zhang A, Rust MJ, Bishop DK. Small Rad51 and Dmc1 Complexes Often Co-occupy Both Ends of a Meiotic DNA Double Strand Break. PLoS Genet. 2015;11(12):e1005653.

14. Fraune J, Schramm S, Alsheimer M, Benavente R. The mammalian synaptonemal complex: protein components, assembly and role in meiotic recombination. Experimental cell research. 2012;318(12):1340–6.

15. de Vries FA, de Boer E, van den Bosch M, Baarends WM, Ooms M, Yuan L, et al. Mouse Sycp1 functions in synaptonemal complex assembly, meiotic recombination, and XY body formation. Genes & development. 2005;19(11):1376–89.

16. Billings T, Sargent EE, Szatkiewicz JP, Leahy N, Kwak IY, Bektassova N, et al. Patterns of recombination activity on mouse chromosome 11 revealed by high resolution mapping. PloS one. 2010;5(12):e15340.

17. de Boer E, Dietrich AJ, Hoog C, Stam P, Heyting C. Meiotic interference among MLH1 foci requires neither an intact axial element structure nor full synapsis. Journal of cell science. 2007;120(Pt 5):731–6.

18. de Boer E, Stam P, Dietrich AJ, Pastink A, Heyting C. Two levels of interference in mouse meiotic recombination. Proceedings of the National Academy of Sciences of the United States of America. 2006;103(25):9607–12.

19. Cole F, Kauppi L, Lange J, Roig I, Wang R, Keeney S, et al. Homeostatic control of recombination is implemented progressively in mouse meiosis. Nature cell biology. 2012;14(4):424–30.

20. Carofiglio F, Inagaki A, de Vries S, Wassenaar E, Schoenmakers S, Vermeulen C, et al. SPO11-Independent DNA Repair Foci and Their Role in Meiotic Silencing. PLoS Genet. 2013;9(6):e1003538.

21. Paul MW, de Gruiter HM, Lin Z, Baarends WM, van Cappellen WA, Houtsmuller AB, et al. SMoLR: visualization and analysis of single-molecule localization microscopy data in R. BMC Bioinformatics. 2019;20(1):30.

22. Boateng KA, Bellani MA, Gregoretti IV, Pratto F, Camerini-Otero RD. Homologous pairing preceding SPO11-mediated double-strand breaks in mice. Developmental cell. 2013;24(2):196–205.

23. Hamer G, Gell K, Kouznetsova A, Novak I, Benavente R, Hoog C. Characterization of a novel meiosis-specific protein within the central element of the synaptonemal complex. Journal of cell science. 2006;119(Pt 19):4025–32.

24. Hamer G, Wang H, Bolcun-Filas E, Cooke HJ, Benavente R, Hoog C. Progression of meiotic recombination requires structural maturation of the central element of the synaptonemal complex. Journal of cell science. 2008;121(Pt 15):2445–51.

25. Schramm S, Fraune J, Naumann R, Hernandez-Hernandez A, Hoog C, Cooke HJ, et al. A novel mouse synaptonemal complex protein is essential for loading of central element proteins, recombination, and fertility. PLoS Genet. 2011;7(5):e1002088.

26. Sanchez H, Paul MW, Grosbart M, van Rossum-Fikkert SE, Lebbink JH, Kanaar R, et al. Architectural plasticity of human BRCA2-RAD51 complexes in DNA break repair. Nucleic acids research. 2017;45(8):4507–18.

27. Haas KT, Lee M, Esposito A, Venkitaraman AR. Single-molecule localization microscopy reveals molecular transactions during RAD51 filament assembly at cellular DNA damage sites. Nucleic acids research. 2018;46(5):2398–416.

28. Mikhaylova M, Cloin BM, Finan K, van den Berg R, Teeuw J, Kijanka MM, et al. Resolving bundled microtubules using anti-tubulin nanobodies. Nature communications. 2015;6:7933.

29. Pleiner T, Bates M, Gorlich D. A toolbox of anti-mouse and anti-rabbit IgG secondary nanobodies. The Journal of cell biology. 2018;217(3):1143–54.

30. Ristic D, Modesti M, van der Heijden T, van Noort J, Dekker C, Kanaar R, et al. Human Rad51 filaments on double- and single-stranded DNA: correlating regular and irregular forms with recombination function. Nucleic Acids Res. 2005;33(10):3292–3302.

31. Lange J, Yamada S, Tischfield SE, Pan J, Kim S, Zhu X, et al. The Landscape of Mouse Meiotic Double-Strand Break Formation, Processing, and Repair. Cell. 2016;167(3):695–708 e16.

32. Hinch AG, Zhang G, Becker PW, Moralli D, Hinch R, Davies B, et al. Factors influencing meiotic recombination revealed by whole-genome sequencing of single sperm. Science. 2019;363(6433).

33. Howard-Till RA, Lukaszewicz A, Loidl J. The recombinases Rad51 and Dmc1 play distinct roles in DNA break repair and recombination partner choice in the meiosis of Tetrahymena. PLoS Genet. 2011;7(3):e1001359.

34. Faieta M, Di Cecca S, de Rooij DG, Luchetti A, Murdocca M, Di Giacomo M, et al. A surge of late-occurring meiotic double-strand breaks rescues synapsis abnormalities in spermatocytes of mice with hypomorphic expression of SPO11. Chromosoma. 2015.

35. Gray S, Allison RM, Garcia V, Goldman AS, Neale MJ. Positive regulation of meiotic DNA double-strand break formation by activation of the DNA damage checkpoint kinase Mec1(ATR). Open biology. 2013;3(7):130019.

36. Kauppi L, Barchi M, Lange J, Baudat F, Jasin M, Keeney S. Numerical constraints and feedback control of double-strand breaks in mouse meiosis. Genes & development. 2013;27(8):873–86.

37. Peters AH, Plug AW, van Vugt MJ, de Boer P. A drying-down technique for the spreading of mammalian meiocytes from the male and female germline. Chromosome research : an international journal on the molecular, supramolecular and evolutionary aspects of chromosome biology. 1997;5(1):66–8.

38. Essers J, Hendriks RW, Wesoly J, Beerens CE, Smit B, Hoeijmakers JH, et al. Analysis of mouse Rad54 expression and its implications for homologous recombination. DNA repair. 2002;1(10):779–93.

39. van de Linde S, Loschberger A, Klein T, Heidbreder M, Wolter S, Heilemann M, et al. Direct stochastic optical reconstruction microscopy with standard fluorescent probes. Nat Protoc. 2011;6(7):991–1009.

40. Schindelin J, Arganda-Carreras I, Frise E, Kaynig V, Longair M, Pietzsch T, et al. Fiji: an open-source platform for biological-image analysis. Nature methods. 2012;9(7):676–82.

